# Self-organized anteroposterior regionalization of early midbrain and hindbrain/spinal cords using micropatterned human embryonic stem cells

**DOI:** 10.1101/2022.07.22.501065

**Authors:** Tianfa Xie, Han Jiang, Lauren E. Brown, ChangHui Pak, Yubing Sun

**Affiliations:** Department of Mechanical and Industrial Engineering, University of Massachusetts, Amherst, Massachusetts 01003, USA; Department of Biomedical Engineering, University of Massachusetts, Amherst, Massachusetts 01003, USA; Department of Biochemistry and Molecular Biology, University of Massachusetts, Amherst, Massachusetts 01003, USA

## Abstract

Anterior-posterior (AP) spatial regionalization is crucial for central nervous system development. Previous studies suggest that morphogen gradients can induce AP patterning in animal and organoid models. While self-organization in early embryogenesis and neurogenesis has been found using geometrically confined microtissues, spontaneously induced AP patterning has not been reported. Here, we show that circularly micropatterned human pluripotent stem cells self-organize into spatially distinct FOXG1-FOXA1+OTX2+ midbrain-like and HOXB4+ hindbrain/spinal cord-like regions. Notably, the tissue then folds inwardly to form a 3D annular structure, maintaining a distinct boundary between OTX2+ and HOXB4+ zones. The reaction-diffusion of BMP/Noggin plays a key role in the mechanism of AP patterning. Our model is validated by its capability to distinguish the teratogenic effects of valproic acid and isotretinoin. Our work suggests a novel regulatory mechanism for AP patterning and provides a tool for fast screening of teratogens.

## Introduction

To fully develop into the central nervous system, the neural tube must first form a pattern consisting of organized neural progenitors along the anterior-posterior (AP) axis (*1*). A series of three vesicles then emerge at the anterior end of the neural tube (*2*). These vesicles are known as the forebrain (prosencephalon, which soon divides into telencephalon and diencephalon), the midbrain (mesencephalon, which grows into the adult brainstem), and the hindbrain (rhombencephalon, which subsequently splits into the rostral metencephalon and caudal myelencephalon) (*2, 3*). The rostral metencephalon then develops into pons and cerebellum while the caudal myelencephalon grows into medulla oblongata (*3*). Secondary organizing centers such as the midbrain-hindbrain boundary, also termed as the isthmus, which functions as an organizer for the midbrain and hindbrain, form subsequently to impose local signaling gradients to sequentially pattern the neural tube into subregions (*1, 4, 5*). Specifically, the midbrain-hindbrain boundary is an evolutionarily conserved structure that forms at the junction of the anterior OTX2^+^ midbrain region and the posterior GBX2^+^ hindbrain region, followed by constriction morphogenesis that separates the adjacent mesencephalon and rostral metencephalon into distinct compartments. Such compartmentalization is essential for proper cell differentiation in later stage development (*6*).

Based on studies (*7–9*) using model organisms such as *Xenopus*, chicks, and zebrafish, it is understood that AP patterning is regulated by several diffusible factors including retinoic acid (RA), sonic hedgehog (SHH), fibroblast growth factors (FGFs), bone morphogenetic proteins (BMPs), and WNTs (*10*). These factors are secreted from surrounding tissues such as the somites, notochord, and dorsal ectoderm to form gradients of signals (*1, 3*).

Recently, *in vitro* development models using human pluripotent stem cells (hPSCs) have provided opportunities for investigating how neural tube is developed and patterned in human (*11–13*). For instance, dorsal-ventral patterning models have been successfully generated via guided self-organization of neural cysts (*14, 15*), by microfluidic diffusion-based gradient generators (*16*), or by organoid fusion techniques (*17, 18*). In addition, by activating WNT or RA signaling, forebrain, midbrain, or rostral hindbrain/spinal cord fates could be obtained in a dose-dependent manner from hPSCs using a defined protocol (*14, 19, 20*). More recently, microfluidic devices have been developed to induce WNT, RA and FGF8 signaling gradients, which generate patterned neural tissue with AP gene expression patterns observed in mouse embryos (*21, 22*).

In addition to external morphogen gradient induced cell fate patterning, we and others have demonstrated that spatially micropatterned hPSCs could serve as unique in vitro systems to study early embryogenesis such as gastrulation (*23–25*), neurulation (*26–30*), and organogenesis including heart (*31*), liver (*32*), and pancreas (*33*). Geometrically confining single or a colony of cells has been achieved by engineering the spatial distribution of extracellular matrix proteins via micro-contact printing with a stamp or via photo-oxidation of a cell-repellent substrate through a photomask (*34, 35*). It has been revealed that micropatterning provides both biomechanical and biochemical cues to guide hPSCs differentiation (*23, 25, 26, 36*). The mechanical stress in cells is relatively higher at the boundary of circular micropatterns and regions with large curvature (*37*). The mechanosensitive activation of BMP and Wnt signals directly regulates cell fate specification (*26, 36*). Moreover, recent studies have shown that the reaction-diffusion (RD) of BMP4 and Noggin leads to spatially patterned BMP activities in micropatterned hPSCs, which explains the gastrulation-like layering in the micropatterned hPSCs (*23, 25, 38*). The two-component RD model, first introduced by Alan Turing in 1952, requires the interaction between an activator and inhibitor pair of morphogens (*39*). Here, the activator, BMP4, induces the expression of itself and its inhibitor, Noggin, while Noggin inhibits the expression of BMP4. The RD of BMP/Noggin thus generates a spatially patterned BMP activity that directly regulates cell fate specification. We further demonstrated that the combinatory effects of the RD of BMP and Wnt regulated the complete ectoderm patterning in confined hPSCs (*29*).

Here, we showed that circularly micropatterned human pluripotent stem cells self-organized into spatially distinct FOXG1-FOXA1+OTX2+ midbrain-like and HOXB4+ hindbrain/spinal cord-like regions by regulating RA, BMP, SHH, and WNT signals. We found that the reaction-diffusion of BMP/Noggin played a role in AP regionalization. We further demonstrated that the model could distinguish the effects of different teratogens on the midbrain and hindbrain/spinal cord development.

## Results

### Generation of AP patterned human neural tissues

We first investigated whether activating RA, SHH, BMP, and WNT signaling pathways could induce anterior and posterior neuroepithelial cells simultaneously from hPSCs monolayers. After two days of neural induction with fully defined Essential 6 (E6) media supplemented with a TGF-β inhibitor SB431542 (SB, 10 µM), we induced H9 human embryonic stem cells (ESCs) with added neural tube morphogenic factors including RA (500 nM), BMP4 recombinant protein (5 ng/ml), SHH pathway agonist (SAG, 1 µM), and WNT pathway agonist CHIR99021 (CHIR, 60 nM) for two or three more days (Figure S1). By continuously inhibiting TGF-β and activating RA, SHH, BMP, and WNT signals, we found that H9 SOX10:: GFP human ESCs could successfully differentiate into both anterior and posterior neural tissues as verified by co-staining HOXB4 as a hindbrain marker and OTX2 as a marker for both posterior forebrain and midbrain progenitors (Figure S2). However, both cell types were randomly distributed and mixed without distinct spatial organization (Figure S2).

As our previous work demonstrated that geometrically confined hPSCs could self-organize into patterned neuroectoderm (*26, 29*), we next studied whether geometrical confinement could facilitate the formation of spatial AP patterning. After plating human ESCs in circular micropatterned substrate (diameter, 600 µm) and inducing cell differentiation using the same protocol (Figure S1), we found that AP patterned human neural tissues were successfully generated. On day 6, human ESCs self-organized into structures with two distinct zones (Figure 1A, top panel). The cells located at the center of the micropattern were OTX2^+^, indicating a forebrain/midbrain cell fate (*5*). Hindbrain/spinal cord marker HOXB4^+^ cells are predominately located at the boundary of the micropattern. A distinct boundary could be observed between the OTX2 and HOXB4 layers, as seen in the epifluorescence images, heatmap, and intensity plots (Figure 1, A, B, and C). We observed similar AP patterning of OTX2 and HOXB4 using a human-induced pluripotent stem cell (iPS) line (*40*) (Figure S3).

**Figure 1.**
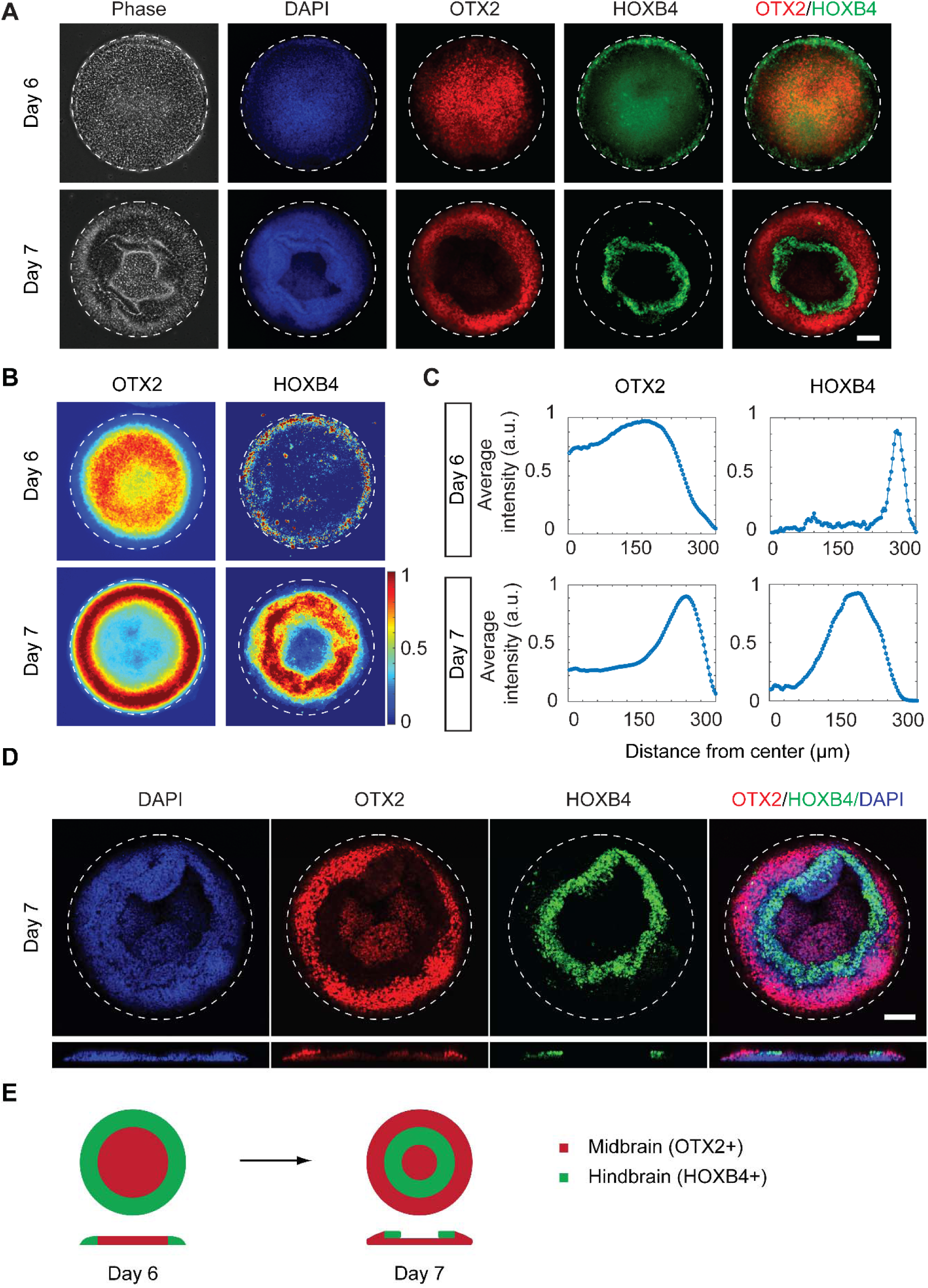
hPSCs self-organized into anteroposterior patterning of midbrain and hindbrain/spinal cord on micropatterned substrates. (**A**) Representative phase-contrast and immunostaining fluorescence images showing the spatially patterned expression of anterior (OTX2) and posterior (HOXB4) neural markers on day 6 and day 7. The OTX2 and HOXB4 merged image was shown. Cell nuclei were counterstained with DAPI. (**B**) Colorimetric maps showing the average fluorescent intensity of OTX2 and HOXB4 staining. The intensity of each pixel of the images was normalized to the maximum value for each image. n > 15. (**C**) Plots showing the quantitative average intensity in relation to the distance to the center of the pattern. (**D**) Representative confocal images and the reconstructed side view showing the three-dimensional architecture and the spatial expression of OTX2+ and HOXB4+ cells on day 7. Cell nuclei were counterstained with DAPI. (**E**) A schematic showing the self-organized anteroposterior patterning generated from human ESCs as well as the folding process and 3D structural change from day 6 to day 7. In these experiments, H9 human ESCs were differentiated with the induction protocol. Scale bar, 100 μm.

Surprisingly, a drastic change in cell positioning was observed after extending the tissue culture. We found that on day 7, HOXB4^+^ cells were located within a ring region in the middle of the micropattern (Figure 1A) and that the boundary cells expressed OTX2. To elucidate the tissue structure, we first performed z-stack confocal microscopy to visualize the cross-section profile of the micropatterned human neural tissue. As revealed in the cross-sectional view, HOXB4^+^ cells stacked on top of the bottom layer of OTX2^+^ cells, suggesting that the HOXB4^+^ cells that were located on the boundary on day 6 subsequently folded inwardly to the center of the micropattern to form the ring structure (Figure 1, D and 1E). We next performed time-lapse live-cell imaging from day 6 to day 7 to capture the dynamics of the process (Figure S4). This morphological change was then confirmed to be caused by cell folding (Figure S4 and movie S1). The bright-field images and the movie showed that the front cells (presumably HOXB4^+^ cells) were moving inwardly while maintaining a distinguishable boundary with the follower cells (presumably OTX2^+^ cells), which is consistent with our immunostaining results.

To further characterize the cell fates, we performed immunostaining for other neural markers including PAX6 (as a neuroepithelial marker), N-cadherin (as a neuroepithelial marker), FOXG1 (as an anterior forebrain marker (*18*)), and GBX2 (as an anterior hindbrain marker (*41*)) (Figure S5). Z-stack confocal microscopy results showed that cells in both layers were PAX6^+^ and N-cadherin^+^ on day 7 (Figure S5), further confirming the neuroepithelial identity of the cells (*20, 26*). In addition, the immunostaining results showed that the FOXG1 signal was not detectable across the micropattern and that very few GBX2 expressing cells colocalized with OTX2^+^ region (Figure S5), suggesting OTX2^+^ midbrain and GBX2^+^ anterior hindbrain were not completely separated at this early stage. To distinguish whether OTX2^+^ cells are posterior forebrain or midbrain cell fates exclusively, and exclude the possibility of choroid plexus fate (*42*), we performed further experiments by immunostaining two additional midbrain markers including engrailed-1 (EN1) and PAX2 (*43*), a choroid plexus marker TTR (transthyretin (*44–46*)), and FOXA1 (*19, 47, 48*) on day 6 (Figure S6-7). FOXA1 has been used as a marker for floor plate cells and midbrain dopaminergic progenitors since it is predominantly expressed in ventral midbrain with an only detectable level in hindbrain but not in forebrain (*19*). The results showed that the cells in the center of the micropattern were TTR^−^ (Figure S7). The cells in the center of the pattern predominantly showed positive FOXA1 staining while the edge of the pattern only showed detectable FOXA1 signal. These results further confirmed the midbrain fate of the cells in the center of the pattern and excluded the possibility of forebrain fates and choroid plexus (Figure S7). In contrast, EN1 and PAX2 were not found across the micropattern (Figure S6). This is possibly because the induction period in our protocol is too short for the expression of those markers. Taken together, the majority of FOXG1^−^ TTR^−^ OTX2^+^ FOXA1^+^ cells in the center of the micropattern have midbrain fate, which are segregated from HOXB4^+^ posterior hindbrain/spinal cord cells in the edge.

### The mechanism of AP patterning of OTX2 and HOXB4 regions

To understand how the OTX2/HOXB4 pattern was formed, we studied three possible known mechanisms of organization, which are cell sorting, differential mechanical stress, and reaction-diffusion (RD) derived biochemical gradient. Firstly, differential tissue interfacial tension has been proposed to explain cell sorting and compartmentalization (*49, 50*). To test whether the AP patterning of OTX2 and HOXB4 was formed via cell sorting, we fixed the cells and performed immunostaining at two earlier time points including 136 h and 140 h (Figure 2A) and compared with the results at 144 h (i.e., day 6). We found that at 136 h, OTX2^+^ cells had emerged in the pattern center while the HOXB4 signals were very weak (Figure 2A). At 140 h, the HOXB4 signals had become detectable at the boundary of the micropatterns (Figure 2A). As the mixture of these two types of cells at an earlier stage was not found, it is unlikely that cell sorting is involved in the formation of two separated layers of midbrain and hindbrain/spinal cord tissues.

**Figure 2.**
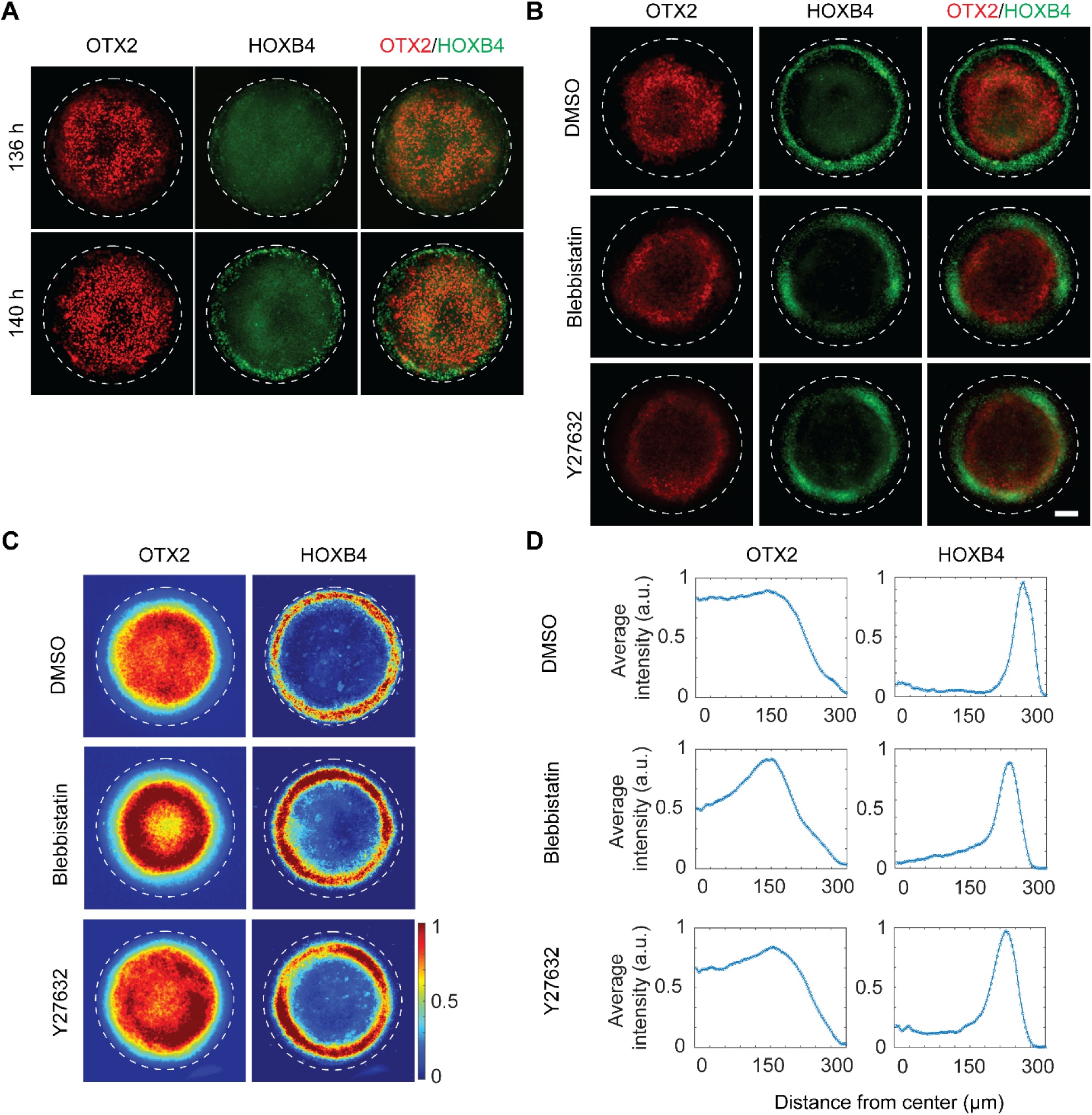
Cell sorting and differential mechanical stress are not the mechanisms that drive the formation of HOXB4 and OTX2 patterning. (**A**) Representative immunostaining fluorescence images showing the spatial expression of neural markers: OTX2 and HOXB4 at 136 hours and 140 hours. The OTX2 and HOXB4 merged image was shown. (**B**) Representative immunostaining fluorescence images showing the spatial expression of OTX2 and HOXB4 for the DMSO control group, Blebbistatin (a myosin inhibitor, 10 µM) treated group, and Y27632 (a ROCK inhibitor, 10 µM) treated group. The OTX2 and HOXB4 merged image was shown. (**C**) Colorimetric maps showing the average fluorescent intensity of OTX2 and HOXB4 staining. The intensity of each pixel of the images was normalized to the maximum value for each image. n > 15. (**D**) Plots showing the quantitative average intensity of OTX2 and HOXB4 in relation to the distance to the center of the pattern. Scale bar, 100 μm.

We and others have reported that differential mechanical tension controls cell fates determination such as neural plate and neural plate border cells (*26*). Thus, we asked whether differential mechanical tension of the boundary and center cells contributed to the AP patterning. We treated the cells with myosin inhibitor (Blebbistatin, 10 µM) or ROCK inhibitor (Y27632, 10 µM), two small molecules widely used to inhibit cellular actomyosin contractility. After treatment with these two drugs from day 4 to day 6, we found that the AP patterning of OTX2 and HOXB4 remained the same as the vehicle control (Figure 2B-D). In addition, we performed immunostaining to investigate whether HOXB4 overlapped with cell nuclei localized Yes-associated protein (YAP), which is a Hippo signaling pathway effector and has been identified as a key mechanotransducer that senses mechanical stimuli. We found that most HOXB4^+^ cells did not show significant overlaps with nuclear YAP^+^ cells (Figure S8). Together, these results suggested that differential mechanical tension was not required for AP patterning.

Our prior work showed that RD of multiple morphogens may synergistically regulate spatial cell fate patterning in micropatterned hPSCs (*29*). To test this possibility, we first investigated the effects of pattern sizes on the regionalization of cell fates since RD patterns changes with tissue size. The immunostaining images showed that the AP patterning remained the same for smaller pattern sizes such as 200 or 400 μm. Interestingly, some colonies with diameters of 800 or 1000 μm showed a periodic regionalization of HOXB4^+^ cells with OTX2^+^ cells flanked by two layers of HOXB4^+^ cells at the boundary and the center of the pattern (Figure 4A). This result suggested that HOXB4 cell fate was determined by one or more morphogens with their activities changing periodically across the pattern, which is a characteristic of RD.

We next sought to elucidate the functional roles of SHH, RA, WNT, and BMP in the AP patterning by treating the cells with inhibitors of these signaling pathways from day 4 to day 6. We found that treating cells with GANT61, a sonic hedgehog pathway inhibitor, did not change the OTX2 and HOXB4 localizations (Figure S9). In contrast, cells treated with AGN193109 (a RA signal inhibitor), IWP2 (a WNT inhibitor), or LDN193189 (a BMP inhibitor) displayed weaker HOXB4 signal and disrupted patterning of HOXB4 and OTX2, suggesting the involvement of RA, WNT and BMP4, but not SHH signaling. To clarify whether their contributions are permissive or instructive, we changed the concentrations of RA, CHIR, and BMP4 added to the media, respectively. We found that lowering RA concentration to 0.1 µM led to weaker HOXB4 signals without changing the relative localization of OTX2 and HOXB4 expressions (Figure 3A, B, and C). Similarly, lowering CHIR concentration to 30 nM or increasing it to 600 nM did not significantly change the AP patterning. Notably, the cells induced with high CHIR concentration (600 nM) on day 6 displayed similar patterns that were observed in cells induced with intermediate CHIR (60 nM) on day 7 (Figure 3D, E, and F and Figure 1A), which can be explained by the effects of WNT activation on cell proliferation. However, when cells were induced with a high concentration of BMP4 (10 ng/ml), the area covered by OTX2^+^ cells became significantly smaller and the peak of HOXB4 signals shifted dramatically inwardly to the center of the micropattern (Figure 4, B, C, and D). Reducing BMP4 concentration to 1 ng/ml led to the expansion of the OTX2^+^ zones. Together, our data suggest that RA and WNT are permissive signals that are required for the caudalization, while the BMP4 signal instructs the AP patterning.

**Figure 3.**
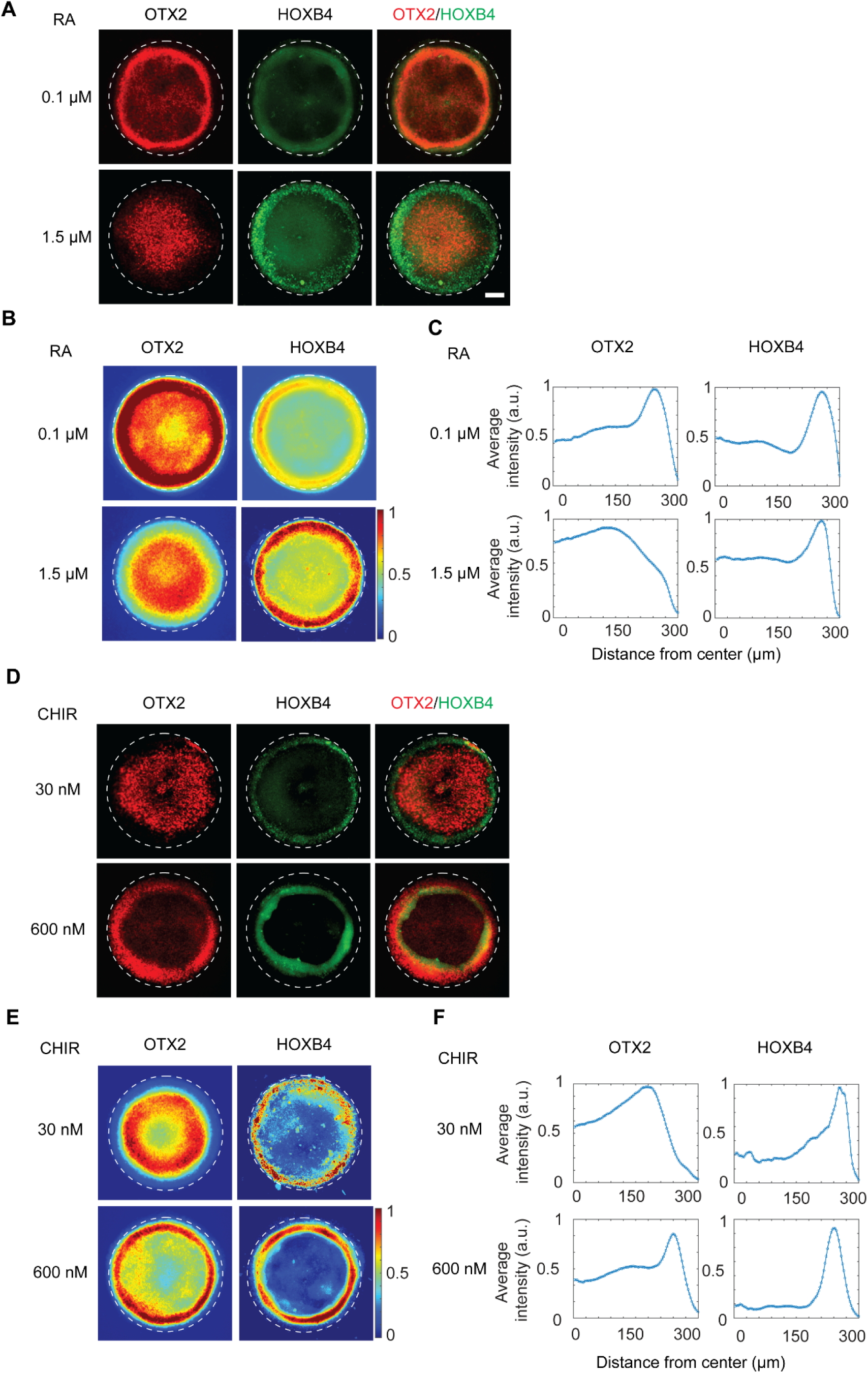
The effects of RA and WNT activation levels on AP patterning. (**A**). Representative immunostaining fluorescence images showing the spatial expression of OTX2 and HOXB4 of human ESCs induced with a lower concentration (0.1 µM) or a higher concentration (1.5 µM) of all-trans retinoic acid (RA) on day 6. (**B**). Colorimetric maps showing the average fluorescent intensity of OTX2 and HOXB4 staining. The intensity of each pixel of the images was normalized to the maximum value for each image. n > 15. (**C**). Plots showing the quantitative average intensity of OTX2 and HOXB4 in relation to the distance to the center of the pattern. (**D**). Representative immunostaining fluorescence images showing the spatial expression of OTX2 and HOXB4 of human ESCs induced with a lower concentration (30 nM) or a higher concentration (600 nM) of WNT activator CHIR99021 on day 6. (**E**). Colorimetric maps showing the average fluorescent intensity of OTX2 and HOXB4 staining. The intensity of each pixel of the images was normalized to the maximum value for each image. n > 15. (**F**). Plots showing the quantitative average intensity of OTX2 and HOXB4 in relation to the distance to the center of the pattern. Scale bar, 100 μm.

**Figure 4.**
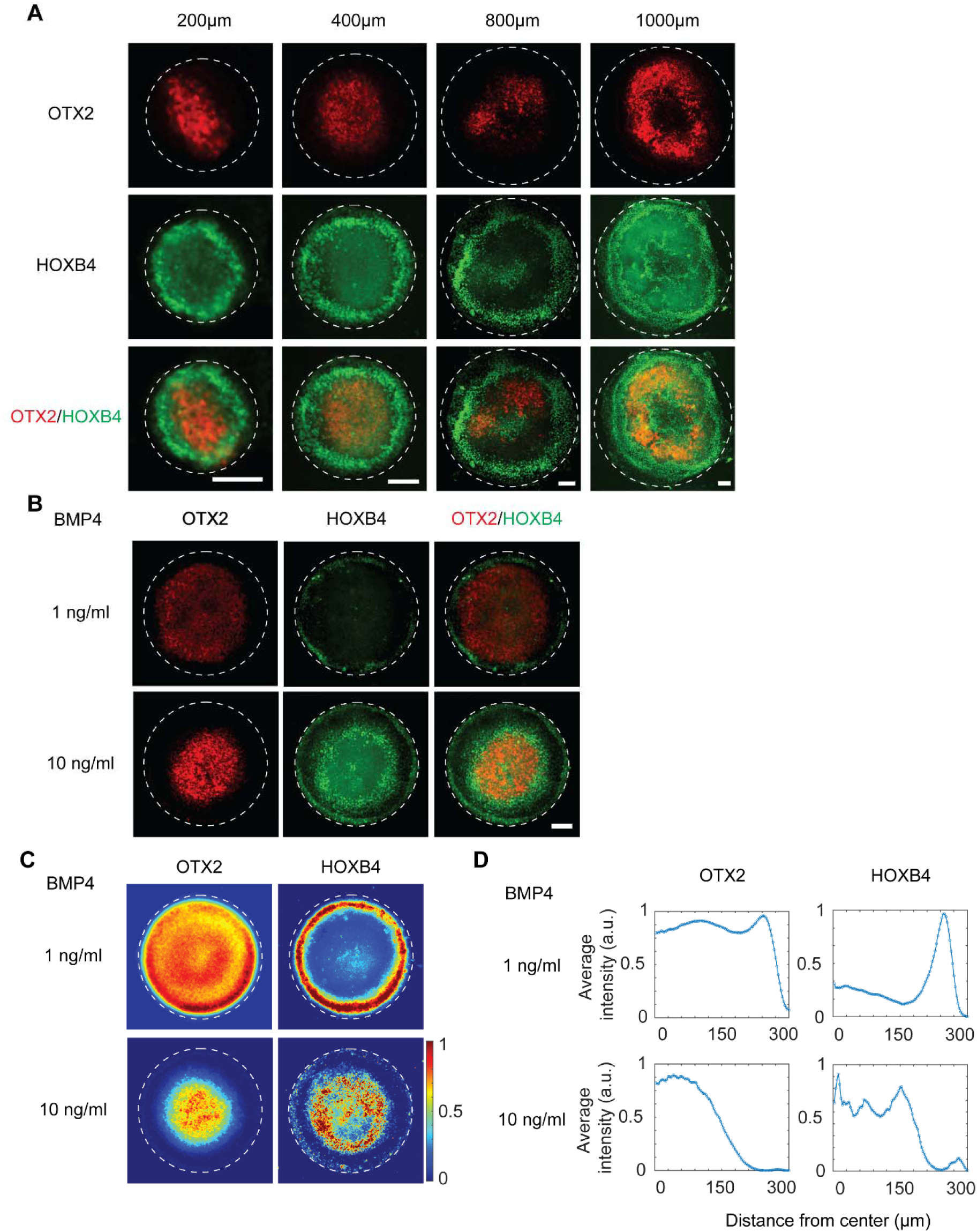
Reaction-diffusion of BMP/Noggin regulates the anteroposterior patterning of OTX2 and HOXB4. (**A**) Representative immunostaining fluorescence images showing the spatial expression of OTX2 and HOXB4 across micropatterns on day 6 with diameters of 200 μm, 400 μm, 800 μm, or 1000 μm. (**B**) Representative immunostaining fluorescence images showing the spatial expression of OTX2 and HOXB4 of human ESCs induced with a lower concentration (1 ng/ml) or a higher concentration (10 ng/ml) of BMP4. (**C**) Colorimetric maps showing the average fluorescent intensity of OTX2 and HOXB4 staining. The intensity of each pixel of the images was normalized to the maximum value for each image. n > 15. (**D**) Plots showing the quantitative average intensity of OTX2 and HOXB4 in relation to the distance to the center of the pattern. Scale bar, 100 μm.

### Drug-induced developmental defects in early midbrain and hindbrain/spinal cord

Several teratogens have been identified that disrupt the midbrain/hindbrain/spinal cord development (*51, 52*). For example, valproic acid (VPA) exposure during embryo development causes neural tube defects and other types of malformations such as smaller midbrain size and an increase in the midline gap of the hindbrain (*53, 54*). It also decreases the proliferation of neuronal progenitor cells in early development (*53*). In addition, recent clinical studies show that exposure to isotretinoin during pregnancy was linked to midbrain/hindbrain development malformations featured as disproportionately long midbrain and dysplasia of the quadrigeminal plate and vermis and disrupted AP axis patterning (*52, 55*).

To further validate our model, we investigated the developmental toxicity caused by VPA and isotretinoin on the midbrain/hindbrain/spinal cord patterning. We induced human ESCs into midbrain and hindbrain/spinal cord cells in the presence of low dosages of VPA and isotretinoin. Penicillin G was used as a non-teratogenic control since it has been shown that it does not disrupt neural tube development at its peak plasma concentration (*30, 56, 57*). First, we performed cell proliferation assay to measure cell viability under treatment with different concentrations of these three tested drugs (Figure S10). Based on the cell viability curves, we determined the 25% inhibition concentrations (IC_25_) for these three drugs to find their range of non-cytotoxic concentration levels (Table S2). Then, we selected three concentrations designated as the low, medium, and high concentrations based on the IC_25_ levels and their peak plasma concentrations (Table S2 and Table S3). To mimic exposure to penicillin G, VPA, or isotretinoin during embryonic development, we treated the human ESCs with these three drugs as well as DMSO as a control group starting from day 2 and performed the same differentiation protocol for midbrain and hindbrain/spinal cord differentiation (Figure S1).

As expected, the results showed that all the low (40 µg/ml), medium (200 µg/ml), and high (1000 µg/ml) dosages of penicillin G treated groups did not alter the midbrain/hindbrain/spinal cord lineage specification and spatial patterning (Figure S11, Figure 5, and Figure S12). In contrast, after treatment with the medium dosage of VPA (0.12 mM), the area covered by OTX2^+^ cells decreased significantly compared to either DMSO control or penicillin G treated group on day 6 while the HOXB4 signal was detectable on day 7 but not on day 6 (Figure 5A and Figure 5G), suggesting that VPA not only inhibited the development of OTX2^+^ midbrain progenitors but also delayed HOXB4^+^ cell fate acquisition. Similar results were shown in the low (0.06 mM) and high (0.25 mM) dosages of VPA-treated groups (Figure S11 and Figure S12). Interestingly, medium concentration (1.63 µM) of isotretinoin-treated colonies showed a randomly mixed population of OTX2^+^ and HOXB4^+^ cells without a boundary (Figure 5A), indicating that isotretinoin disrupted the normal AP patterning of OTX2 and HOXB4. In addition, the results showed that the percentage of OTX2^+^ covered area decreased significantly, compared with the DMSO control or penicillin G treated group (Figure 5A and Figure 5G). The low (1.08 µM) and high (3.25 µM) dosages of isotretinoin treated group displayed similar results (Figure S11, and Figure S12). These observations are consistent with the effects of VPA and isotretinoin on midbrain/hindbrain/spinal cord development *in vivo*, suggesting that our model can distinguish teratogens with different mechanisms (Figure 6).

**Figure 5.**
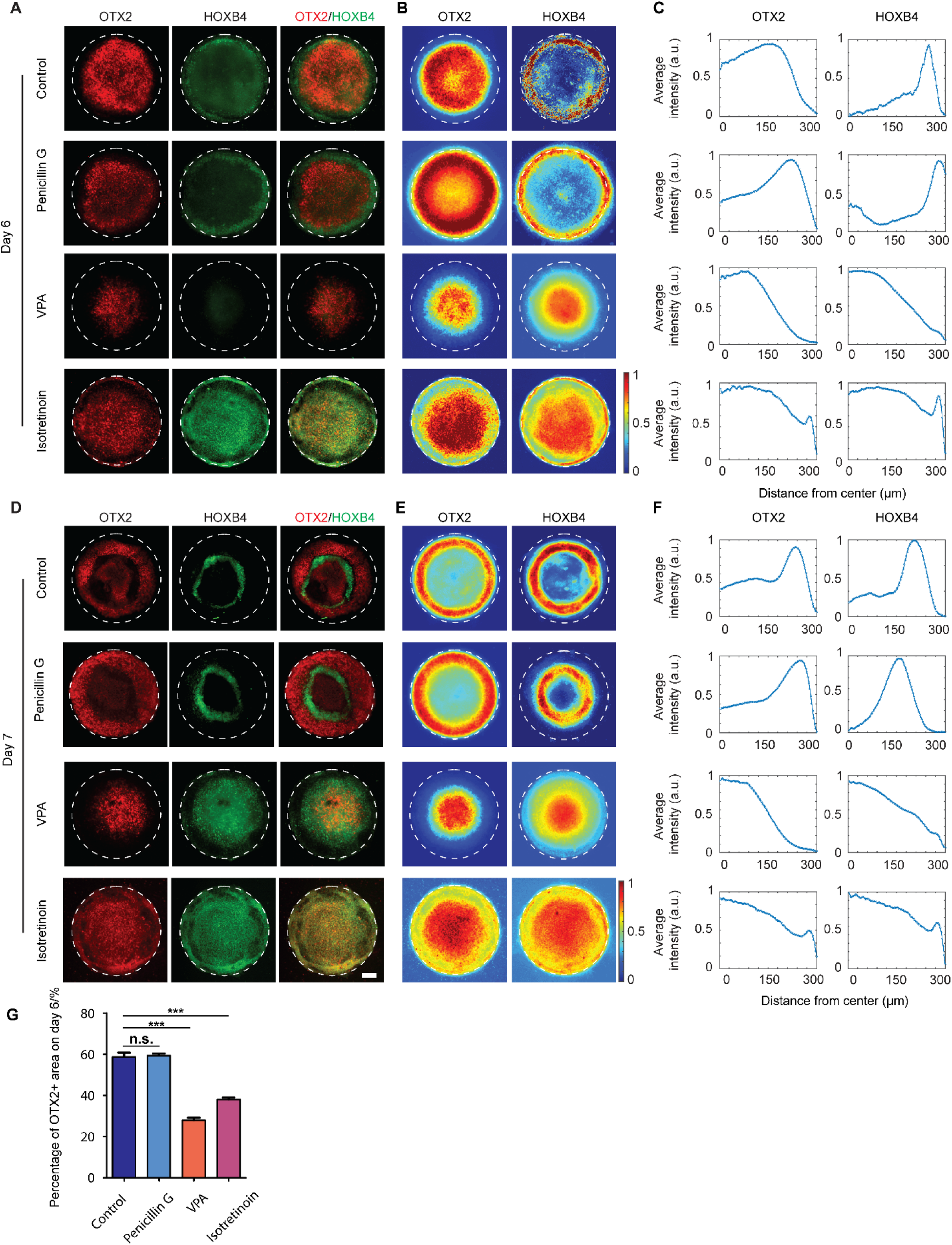
The effects of medium dosages of penicillin G, valproic acid, and isotretinoin on anteroposterior patterning of OTX2 and HOXB4. (**A**) Representative immunostaining fluorescence images showing the spatial expression of OTX2 and HOXB4 in induced human ESCs treated with DMSO (control) and medium dosages of penicillin G, valproic acid, and isotretinoin on day 6. (**B**) Colorimetric maps showing the average fluorescent intensity of OTX2 and HOXB4 staining. The intensity of each pixel of the images was normalized to the maximum value for each image. n > 15. (**C**) Plots showing the quantitative average intensity in relation to the distance to the center of the pattern. (**D**) Representative immunostaining fluorescence images showing the spatial expression of OTX2 and HOXB4 in induced human ESCs treated with DMSO (control) and medium dosages of penicillin G, valproic acid, and isotretinoin on day 7. (**E**) Colorimetric maps showing the average fluorescent intensity of OTX2 and HOXB4 staining. The intensity of each pixel of the images was normalized to the maximum value for each image. n > 15. (**F**) Plots showing the quantitative average intensity in relation to the distance to the center of the pattern. (**G**) The percentage of area covered by OTX2+ midbrain in DMSO control, penicillin G, VPA, and isotretinoin treated groups on day 6. Data are represented as mean ± s.e.m. n.s. (no significance), P > 0.05;***, P < 0.001. Scale bar, 100 μm.

**Figure 6.**
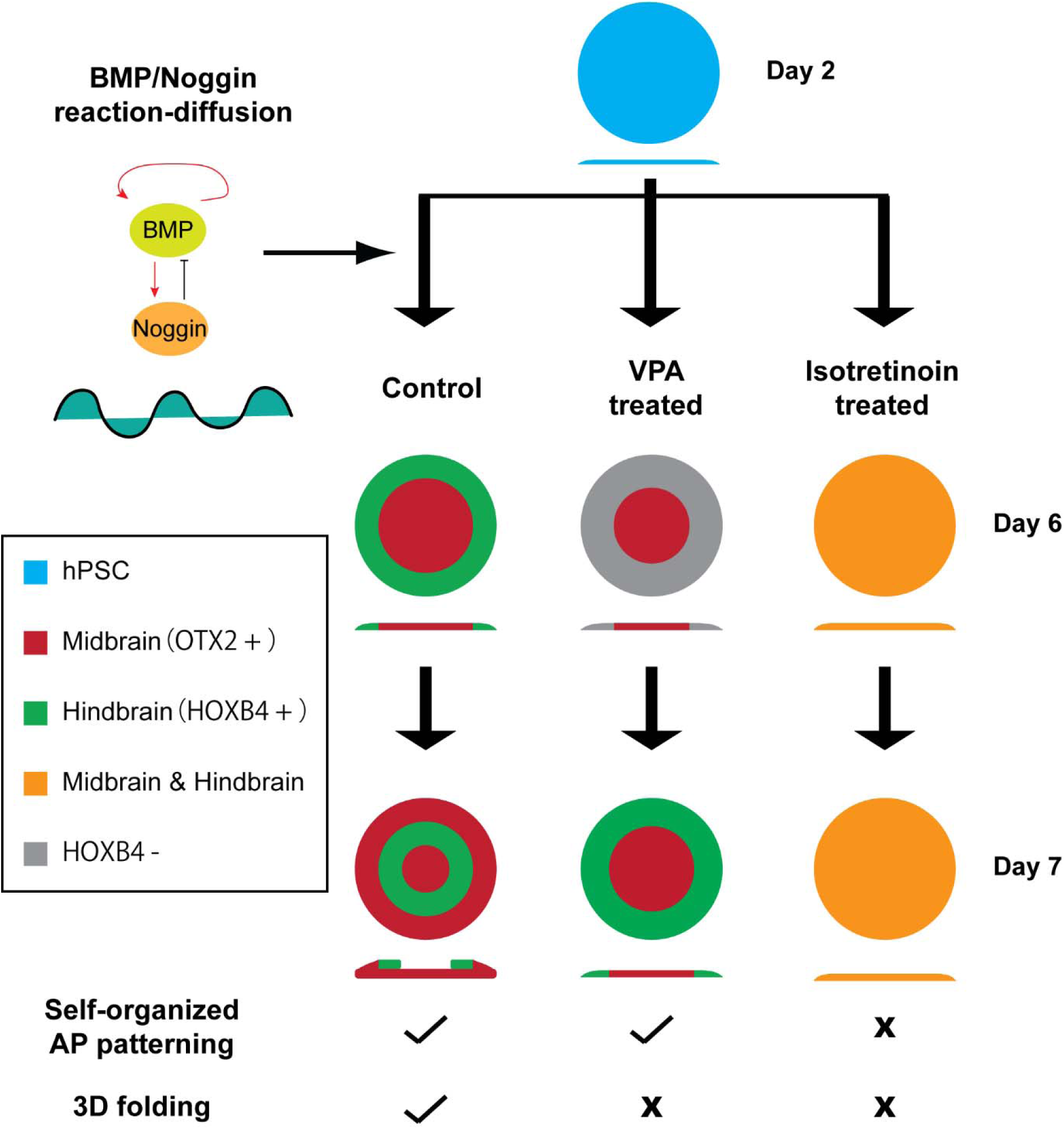
A schematic depicts an *in vitro* model demonstrating self-organized patterning between midbrain and hindbrain/spinal cord cell fates, highlighting its potential in drug testing applications. The model features circularly micropatterned human pluripotent stem cells (hPSCs) that, by day 6, self-organize into distinct zones. These zones exhibit anterior-posterior (AP) patterning of FOXG1-FOXA1+OTX2+ ventral midbrain and HOXB4+ hindbrain/spinal cord fates. The tissue then undergoes inward folding to form a three-dimensional (3D) annular structure while maintaining a clear boundary between the OTX2+ and HOXB4+ zones. The reaction-diffusion of BMP/Noggin plays a key role in the mechanism of midbrain and hindbrain/spinal cord fate patterning. This model can be potentially used in drug testing, which is demonstrated by its capability to differentiate the teratogenic effects of drugs like valproic acid (VPA) and isotretinoin. VPA treatment impairs the development of both midbrain and hindbrain/spinal cord fates, delays anterior-posterior patterning formation on day 6, and leads to 3D folding failure on day 7. Interestingly, isotretinoin treatment leads to a random mix of OTX2+ and HOXB4+ cells, obliterating the anterior-posterior patterning and the formation of 3D structures. Compared to 2D cell culture, the unique phenotypes including completely separated midbrain/hindbrain/spinal cord zones and tissue folding can be used to quantify developmental defects. This innovative model offers an excellent experimental platform for exploring human brain development mechanisms and enhancing drug testing methodologies.

## Discussion

In this work, we demonstrate that micropatterned hPSCs can self-organize into distinct zones with midbrain and hindbrain/spinal cord cell fates, when RA, BMP4, and WNT signals are activated (Figure 6). Micropatterning or geometrical confinement clearly plays an important role in AP pattern formation in our model because micropatterned but not unpatterned hPSCs can form AP patterned cell fates (Figure S1 and Figure 1). While more accurate identification of cell fate is challenging at this early stage and may require additional characterization such as spatial transcriptomics, our immunocytochemistry results reveal the segregation of two cell populations with midbrain fate and posterior hindbrain/spinal cord fate, respectively. The formation of a definitive midbrain-hindbrain boundary may require additional morphogens such as FGF8 to specify midbrain and anterior hindbrain fates. To investigate how micropatterning facilitates the formation of spatial AP patterning, we studied the role of mechanical stress and biochemical cues since geometrical confinement controls the distribution of both factors.

It has been demonstrated that differential mechanical tension across micropatterns controls cell fate determination such as neural plate and neural plate border cells (36), and mesoderm specification (48). In our model, we found that differential mechanical tension is not required for AP patterning because cells can still form AP patterns with inhibited actomyosin contractility (Figure 2B-D and Figure 6). Our mechanistic studies support that BMP/Noggin reaction-diffusion dictates the AP patterning (Figure 6). We found that two layers of HOXB4+ cells can be generated in the same pattern, which is a characteristic of RD (Figure 4A). In addition, OTX2+ cells layer thickness and the peak of HOXB4 signals can be tuned by changing the BMP4 concentrations (Figure 4, B, C, and D). Our findings that the concentration of BMP4 but not CHIR regulated the sizes of OTX2^+^ zones and location of HOXB4^+^ zones were in contrast with findings that WNT signal dictated AP patterning in neural tube-like tissues (*21*). One possibility is that BMPs crosstalk with WNT signals. For instance, it has been revealed that BMP4 could trigger the activation of WNT signaling pathway during gastrulation (*58, 59*). Moreover, it has been found that BMP4 and WNT activations led to midbrain fate in human brain organoids (*60*) and BMP activities inhibited hindbrain fates in *Xenopus laevis* (*61*). Further studies will be needed to address the detailed mechanism of how BMP activity regulates the HOXB4 or OTX2 cell fate determination and whether it is via the WNT and/or RA pathway.

VPA has been known to be teratogenic since the early 1980s (*62*) and exposure to VPA during human embryo development has been found tightly linked to neural tube defects (*51*) and malformations in the midbrain and hindbrain (*53, 54*), as well as neurological disorders such as severe cognitive defects, autism spectrum disorder, and schizophrenia (*63*). Recently, cytoskeletal protein MARCKSL1, which is an actin-stabilizing protein essential for dendrite morphogenesis and synapse maturation, has been identified to mediate VPA-induced morphological defects in developing neurons (*64*). In addition, it has been found with a human cortical organoid-on-a-chip model that the transcriptional regulation of genes such as KLHL1, LHX9, and MGARP could be a mechanism for prenatal VPA exposure induced postnatal brain disorders such as autism (*65*). Isotretinoin exposure during early pregnancy has been known to be associated with an increased risk for infant brain malformation (*55, 66*) such as hypoplasia of the cerebellar vermis, microencephaly, vermis hypoplasia, dysplastic quadrigeminal plate, disproportionate size between midbrain and pons (*55*). Known as a retinoic acid analog, isotretinoin (13-cis retinoic acid) likely acts on the retinoic acid signaling pathway to regulate the expression of *Hox* genes development and thus the spatial patterning of the midbrain and hindbrain/spinal cord. However, the detailed mechanism of these two drugs causing developmental defects, especially in the human genetic background, remains to be investigated. Using our model, it is promising to investigate how VPA and isotretinoin cause defects in human early midbrain and hindbrain/spinal cord development and patterning (Figure 6).

It is notable that our *in vitro* midbrain/hindbrain/spinal cord model so far can mimic only the early-stage development of the human brain since the total duration of organized cell culture is limited due to the geometrical confinement. Extended culture of cells beyond 7 days leads to disorganized cell aggregates or clusters. Despite these disadvantages, our model is complementary to current animal models and traditional 2D cell culture systems. Compared to animal models, our system utilizes pluripotent stem cells with human genetic background and features simpler experimental procedures and a shorter time frame (∼ 2 weeks). Compared to 2D cell culture, the unique phenotypes including completely separated midbrain/hindbrain/spinal cord zones and tissue folding can be used to quantify developmental defects. When combined with genetic engineering and automated image processing tools, we envision that our system has the potential to evolve into a high-throughput screening platform for teratogens and toxicology testing.

In conclusion, we have developed an *in vitro* model for AP patterned midbrain/hindbrain/spinal cord tissues using hPSCs by modulating RA, BMP, SHH, and WNT signals. The regionalization of midbrain and hindbrain/spinal cord cells is mainly mediated by the reaction-diffusion of BMP/Noggin. In addition, we show that our model can be used to predict the distinct effects of different teratogens on midbrain/hindbrain/spinal cord development. This model will provide an excellent experimental platform to investigate the mechanism of human brain development and patterning.

## Materials and Methods

### Cell culture

Both SOX10:: EGFP bacterial artificial chromosome hES cell reporter line (H9; WA09, WiCell; NIH registration number: 0062; female; generated by Dr. Lorenz Studer’s lab (*67*)) and control human induced pluripotent stem (hiPS) cell line (male; generated from human primary T-cells using episomal reprogramming method) were cultured in Essential 8 growth medium on hES cell-qualified Geltrex (Thermo Fisher Scientific). Vitronectin, which is known to support the stemness maintenance of hPSCs, was used for both micropatterned and unpatterned hPSC experiments to promote cell adhesion. Karyotyping and mycoplasma testing were performed routinely to ensure genomic integrity. After treatment with EDTA dissociation buffer (5 min at 37 °C), the cells were collected using a cell scraper (BD Biosciences), centrifuged (200 g for 5 min), and re-dispersed in Essential 8 growth medium supplemented with ROCKi (Y27632, 10 µM; Cayman Chemical) before cell seeding. The induction media are comprised of Essential 6 basal media, TGF-β inhibitor SB431542 (SB, Cayman Chemical; 10 µM), WNT agonist CHIR99021 (CHIR, Cayman Chemical; 60 nM), all-trans retinoic acid (RA, Cayman Chemical; 500 nM), smoothened agonist (SAG, Cayman Chemical; 1 µM), and BMP4 recombinant protein (Gibco; 5 ng/mL). The media was half-changed every day during the differentiation process. All cells were cultured at 37 °C and 5% CO_2_.

### Microcontact printing

Soft lithography was used to generate micropatterned polydimethylsiloxane (PDMS) stamps from negative SU8 molds that were fabricated using photolithography. These PDMS stamps were used to generate micropatterned cell colonies using microcontact printing, as described previously (*68*). Briefly, to generate micropatterned cell colonies on flat PDMS surfaces, round glass coverslips (diameter = 25 mm, Fisher Scientific) were spin-coated (Spin Coater; Laurell Technologies) with a thin layer of PDMS prepolymer comprising of PDMS base monomer and curing agent (10:1 *w*/*w*; Sylgard 184, Dow-Corning). PDMS coating layer was then thermally cured at 110 °C for at least 24 h. In parallel, PDMS stamps were incubated with a vitronectin solution (20 µg ml^−1^, in deionized water) for 1 h at room temperature before being blown dry with a stream of nitrogen. Excess vitronectin was then washed away by distilled water and the stamps were dried under nitrogen. Vitronectin-coated PDMS stamps were then placed on top of ultraviolet ozone-treated PDMS (7 min, UV-ozone cleaner; Jetlight) on coverslips with a conformal contact. The stamps were pressed gently to facilitate the transfer of vitronectin to PDMS-coated coverslips. After removing stamps, coverslips were disinfected by submerging in 70% ethanol. Protein adsorption to PDMS surfaces without printed vitronectin was prevented by incubating coverslips in 0.2% Pluronic F127 solution (P2443-250G, Sigma) for 30 min at room temperature. Coverslips were rinsed with PBS before being placed into tissue culture plates for cell seeding. For micropatterned cell colonies, PDMS stamps containing circular patterns with diameters of 600 µm were used if not specifically indicated.

### Immunocytochemistry

4% paraformaldehyde (Electron Microscopy Sciences) was used for cell fixation before permeabilization with 0.1% Triton X-100 (Fisher Scientific) at 4% serum with 0.1% Triton X-100 on an orbital shaker (20 rpm) for 24 hours at 4 °C. Then samples were incubated with primary antibodies (listed in the Table S1) diluted in antibody dilution buffer (4% DS, 0.1% Triton-X 100 in sterile PBS) at 4 °C for 48 hours. Finally, samples were incubated with secondary antibodies at 4 °C for 24 hours. Samples were counterstained with 4,6-diamidino-2-phenylindole (DAPI; Invitrogen) to visualize the cell nucleus. The primary antibodies used in this study include HoxB4 (rat, DSHB, I12, 1:20), OTX2 (goat, R&D Systems, 1:100), and PAX6 antibodies (mouse, Abcam, Ab78545, 1:100). The I12 anti-HoxB4 monoclonal antibody193, developed by Gould, A / Krumlauf, R in MRC National Institute for Medical Research, was obtained from the Developmental Studies Hybridoma Bank (DSHB), created by the NICHD of the NIH and maintained at The University of Iowa, Department of Biology, Iowa City, IA 52242. For immunolabelling, donkey-anti goat IgG-Alexa Fluor 647 (1:1000, Life Technologies), donkey-anti rat IgG-Alexa Fluor 488 (1:1000, Life Technologies), donkey-anti rabbit IgG-Alexa Fluor 555 (1:1000, Life Technologies) secondary antibodies were used.

### Image acquisition and analysis

Confocal images were collected with NIS-Elements AR software using Nikon A1 Resonant scanning confocal inverted microscope. Alexa Fluor 488 was excited with a 488 nm laser line. Alexa Fluor 555 was excited with a 561 nm laser line. Alexa Fluor 647 was excited with a 640 nm laser line. 10× objective was used. Data analysis was performed using the NIS-Elements AR analysis software. All other phase contrast and fluorescence images of micropatterned cell colonies were recorded using an inverted epifluorescence microscope (Leica DMi8; Leica Microsystems) equipped with a monochrome charge-coupled device (CCD) camera. Images were first cropped using a custom-developed MATLAB program (MathWorks; https://www.mathworks.com/) to a uniform square with pattern centroids aligned. Cell colonies with a non-radially symmetric cell density pattern or with disorganized cellular structures were excluded from data analysis. The fluorescence intensity of each pixel in cropped images was normalized by the maximum intensity identified in each image. These normalized images were stacked together to obtain average intensity maps. To plot average intensity as a function of distance from colony centroid, average intensity maps for circular cell colonies were divided into 100 concentric zones with equal widths. The average pixel intensity in each concentric zone was calculated and plotted against the mean distance of the concentric zone from the colony centroid.

### Pharmaceutical inhibitions assays

Blebbistatin (10 µM, Cayman Chemical), Y27632 (10 µM, Cayman Chemical), IWP2 (3 µM, Cayman Chemical), GANT61 (10 µM, Cayman Chemical), LDN193189 (500 nM, Cayman Chemical), and AGN193109 (100 nM, MedChemExpress) were dissolved in dimethyl sulfoxide (DMSO, Sigma-Aldrich). Cells were treated with these drugs for 48 and/or 72 hours at 37 °C.

### Cell proliferation assay (MTS)

To determine the cytotoxicity of drugs, 10,000 hESCs/well were seeded into each well of 96-well plates with E8 supplemented with Y27632 and cultured overnight. The medium supplemented with Y27632 was changed to E8 and then the cells were treated with the selected drugs with 8 serial concentrations with 4 (for isotretinoin) or 5 (for valproic acid and penicillin G) times dilution factor along with solvent for 48 hours. The medium supplemented with drugs was changed daily. The cell proliferation assay was conducted with CellTiter 96® AQ_ueous_ One Solution Cell Proliferation Assay (MTS) reagent (Promega, Cat.No. G3582) following the instruction. The absorbance was measured using a plate reader (Cytation 3 microplate reader, BioTek Instruments Inc, Winooski, VT, USA). The corrected absorbance at 490 nm was plotted versus the concentrations of the drugs. The IC_25_ values were determined manually using GraphPad Prism 7.

### Drug-screening assay

Valproic acid (Cat. No. P4543, Sigma-Aldrich) and penicillin G (Cat. No. 13752-1G-F, Sigma-Aldrich) were dissolved in Milli-Q water. Isotretinoin (Cat. No. 21648, Cayman Chemical) was dissolved in DMSO. The cells were induced with Essential 6 medium supplemented with these drugs as well as other factors starting from day 2 until day 6 and/or 7. The medium was half-changed daily.

### Quantification and statistical analysis

Data are represented as mean ± s.e.m. All the micropatterning experiments were repeated at least three times independently (biological replicates), and in each experiment, at least two technical replicates (replicates within each experiment) were used. Sample sizes were determined based on our previous experience and similar studies of other groups. Samples were randomly allocated into different experimental groups. Investigators were not blinded to group allocation, as no animal/human studies were conducted in this manuscript. Due to intrinsic inhomogeneous cell seeding, patterns with multilayered cellular structures or uneven cell density would inevitably appear in micropatterned cell colonies. Thus, these patterns were manually excluded from image analysis. The number of colonies included in each analysis (N) is mentioned in the figure legends. Statistical analysis was performed using two-sided unpaired Student’s t-test. ***, P < 0.001, **, P < 0.01, *, P < 0.05.

## Acknowledgments

We thank the Conte Nanotechnology Cleanroom Lab for support in microfabrication. We also acknowledge the Light Microscopy Facility where confocal microscopy data were collected. Y.S. was funded by National Science Foundation grant CMMI 1846866 and CBET-2326703, National Institute of Diabetes and Digestive and Kidney Diseases R01DK129990 and National Institute of Mental Health R21MH130843.

## Author Contributions

T.X. and Y.S. designed research, T.X. and H.J. performed experiments, T.X., H.J., and L.B analyzed data, C. P. generated the human induced pluripotent stem cell line. All authors wrote and approved the manuscript.

## Data and materials availability

All data needed to evaluate the conclusions are available in the main text or the supplementary materials. The MATLAB scripts (MicropatternImageAnalysis-V1.0) used in the analysis of micropatterned immunostaining images are made available to the public (Zenodo. https://doi.org/10.5281/zenodo.7450143). All materials used in this work are commercially available. There is no restriction on the availability of any materials used in this work.

