## Supplementary Figures and Tables for "Self-organized anteroposterior regionalization of early midbrain and hindbrain/spinal cords using micropatterned human embryonic stem cells"

**Affiliations**

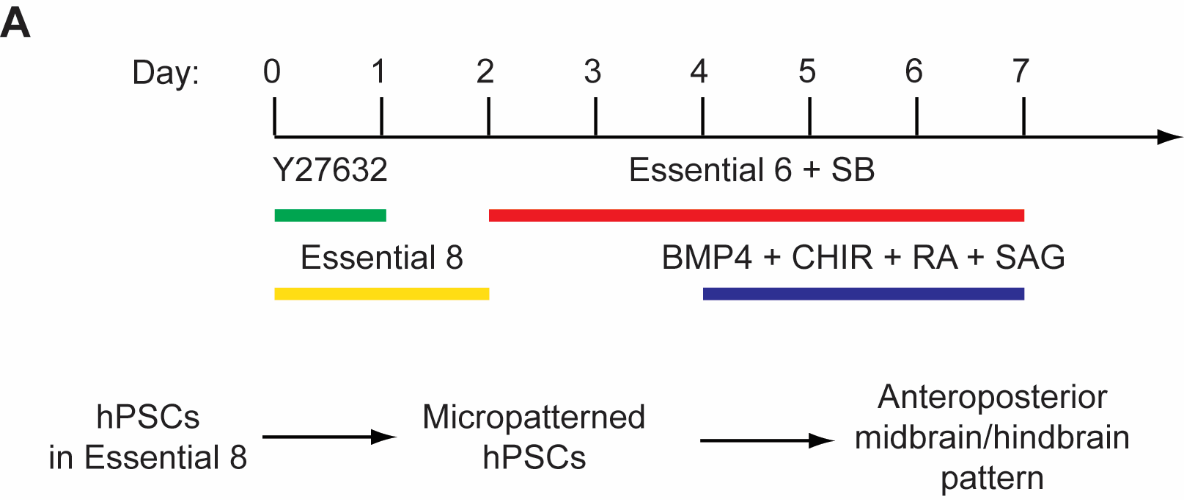

**Figure S1. A schematic showing the culture conditions and induction timeline for anteroposterior midbrain/hindbrain micropatterning**. hPSCs were cultured in Essential 8 medium and then plated onto micropatterned substrates with ROCK inhibitor (Y27632, 10 µM) in Essential 8 medium for one day. The culture medium was changed back to Essential 8 on day 1. Cells were then induced with TGF-β inhibitor SB431542 (SB, 10 µM) in Essential 6 medium starting from day 2. BMP4 recombinant protein (BMP4, 5 ng/ml), CHIR99021 (CHIR, 60 nM), all-trans retinoic acid (RA, 500 nM), and smoothened agonist (SAG, 1 µM) were added to the induction media from day 4 to day 7. The medium was half-changed every day during the induction process. Cells were fixed on day 6 or day 7.

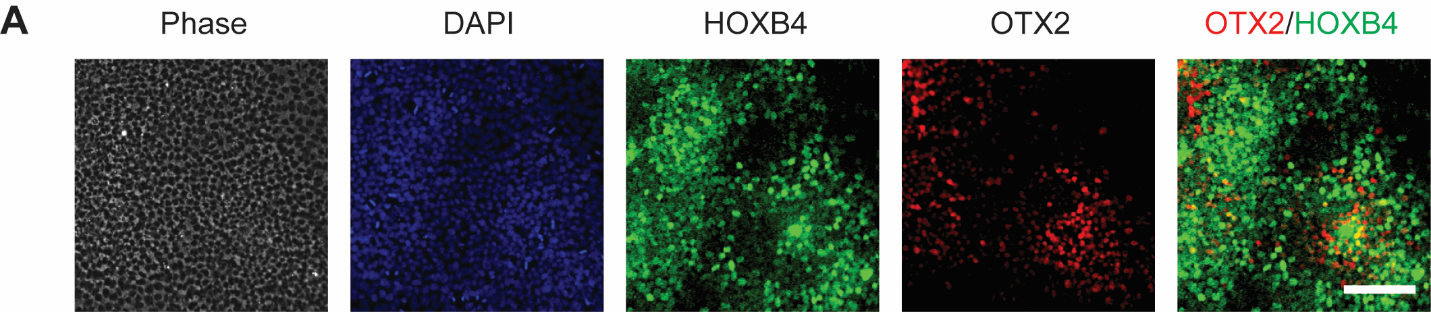

**Figure S2. Induction of anterior OTX2 and posterior HOXB4 cell fates from human ESCs monolayer.** Representative phase-contrast and immunostaining fluorescence images showing the expression of anterior (OTX2) and posterior (HOXB4) neural markers on day 7. Cell nuclei were counterstained with DAPI. Scale bar, 100 μm.

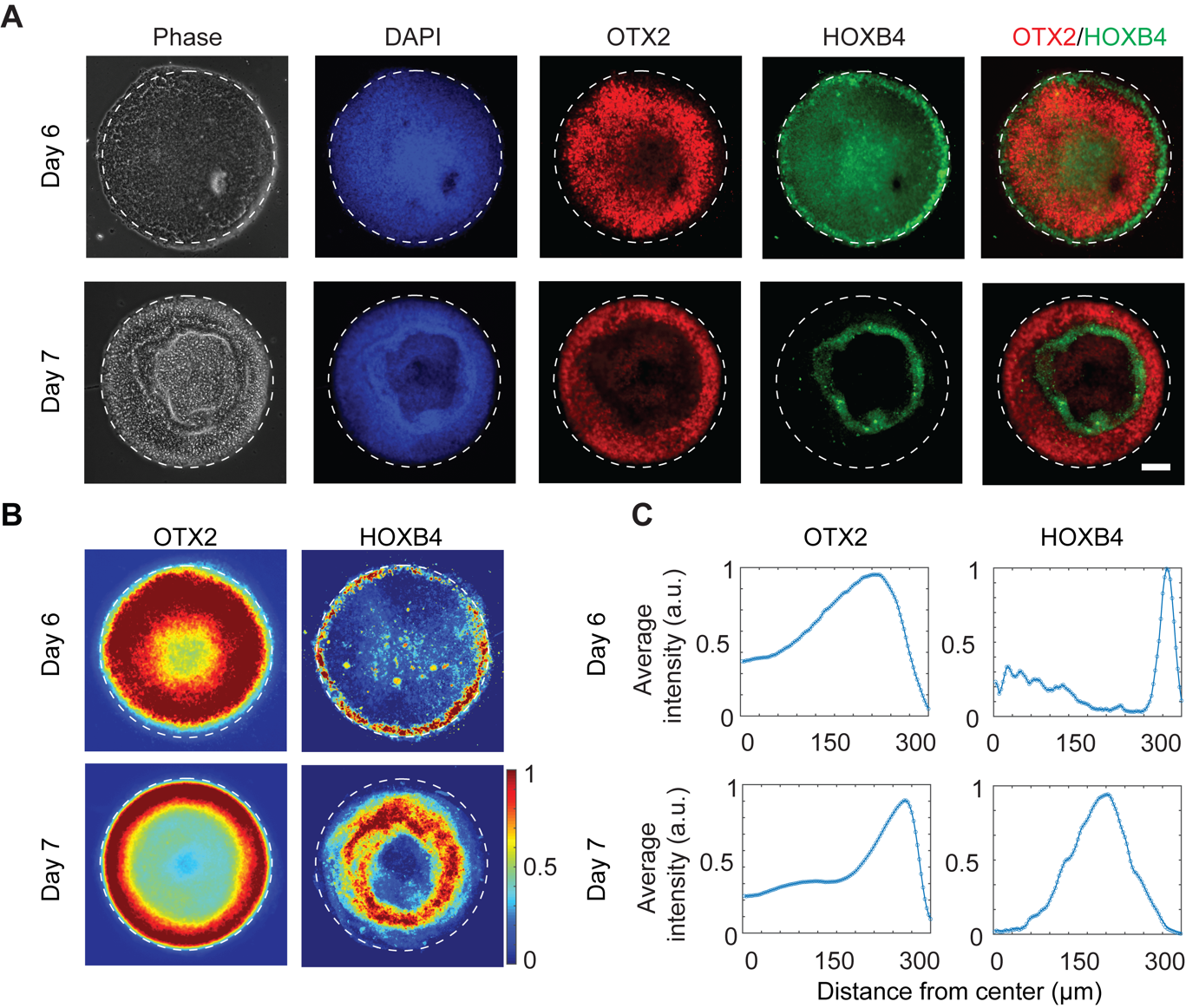

**Fig. S3. hPSCs self-organized into anteroposterior patterning of midbrain and hindbrain on micropatterned substrates.** (A). Representative phase-contrast and immunostaining fluorescence images showing the spatially patterned expression of anterior (OTX2) and posterior (HOXB4) neural markers on day 6 and day 7. The OTX2 and HOXB4 merged image was shown. Cell nuclei were counterstained with DAPI. (B). Colorimetric maps showing the average fluorescent intensity of OTX2 and HOXB4 staining. The intensity of each pixel of the images was normalized to the maximum value for each image. n > 15. (C). Plots showing the quantitative average intensity in relation to the distance to the center of the pattern. In these experiments, the human iPS cells were differentiated with the induction protocol. Scale bar, 100 μm.

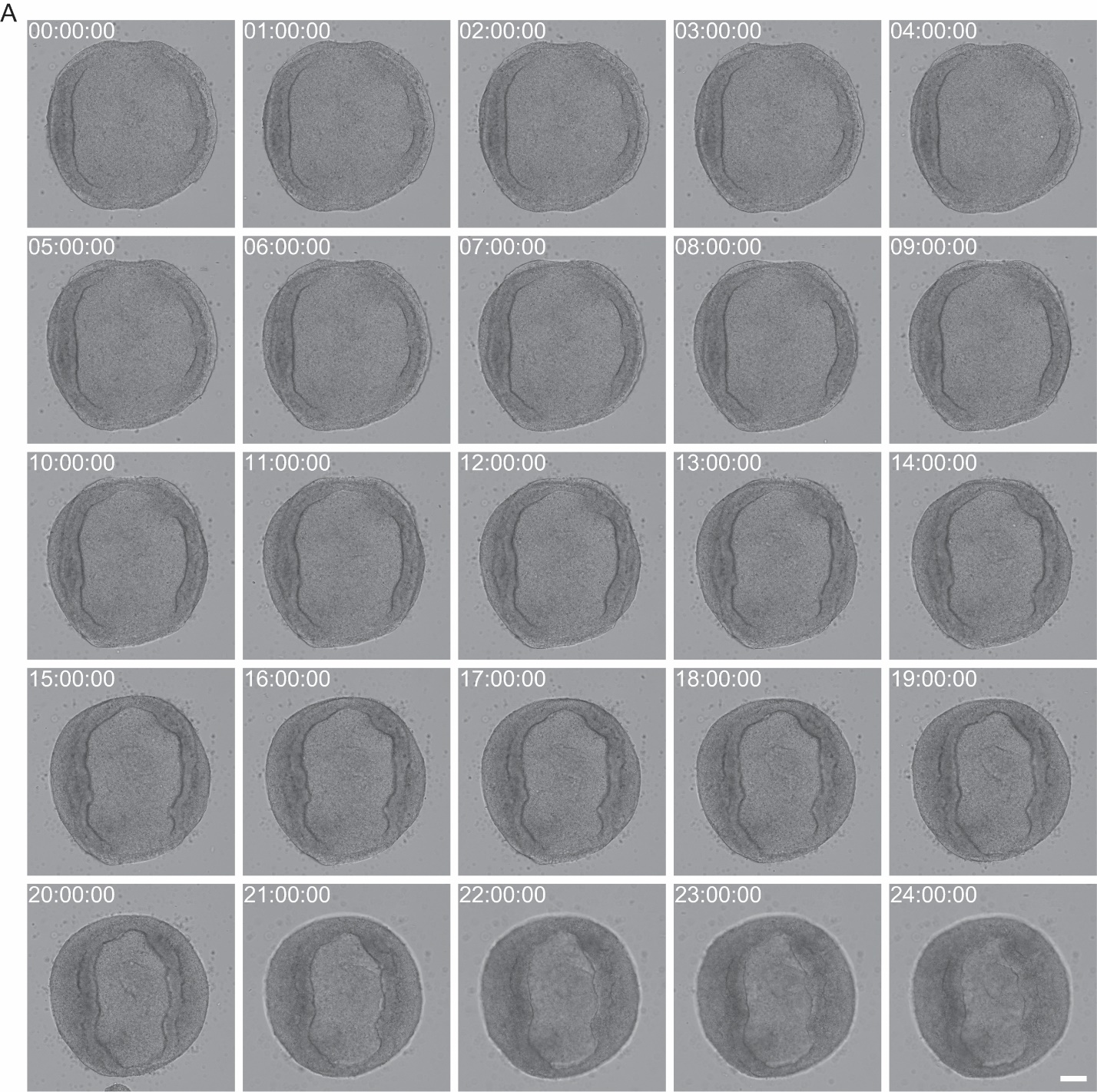

**Figure S4. Time-lapse bright-field images showing the folding process of micropatterned cells from day 6 to day 7.** Micropatterned human ESCs were differentiated with the induction protocol. From day 6 to day 7, the cells were folded inwardly and then formed a multilayer structure in 3D. Scale bar, 100 µm.

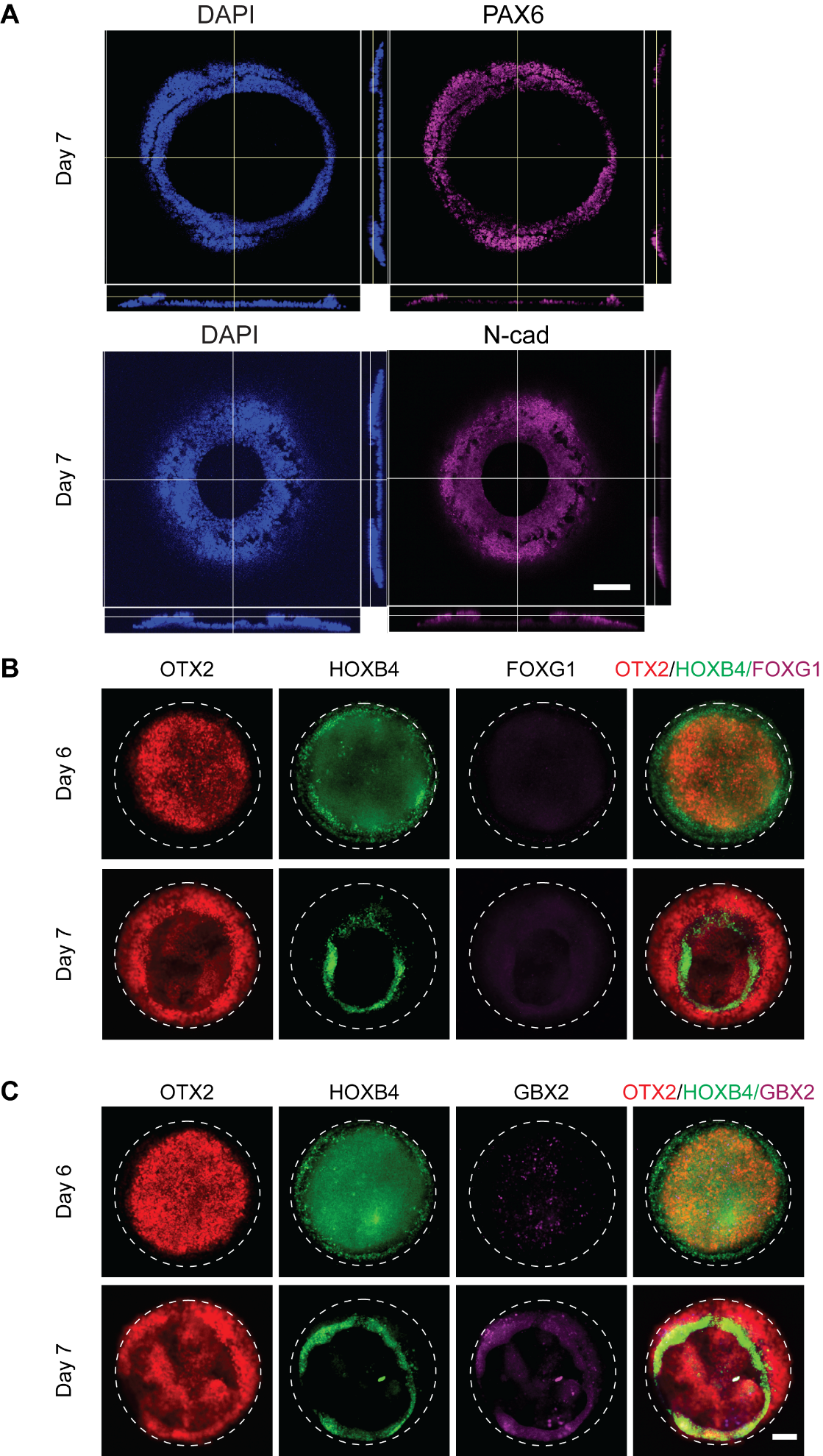

**Figure S5. Characterization of cell fates in self-organized anteroposterior patterned human neural tissue.** (A). Confocal images and the reconstructed side views showing the spatial expression of neuroepithelial markers PAX6 and N-cadherin on day 7. (B). Representative immunostaining fluorescence images showing the spatial expression of neural markers: forebrain and midbrain marker OTX2, hindbrain marker HOXB4, and forebrain marker FOXG1 on day 6 and day 7. (C). Representative immunostaining fluorescence images showing the spatial expression of neural markers: forebrain and midbrain marker OTX2, hindbrain marker HOXB4, and anterior hindbrain marker GBX2 on day 6 and day 7. Scale bar, 100 μm.

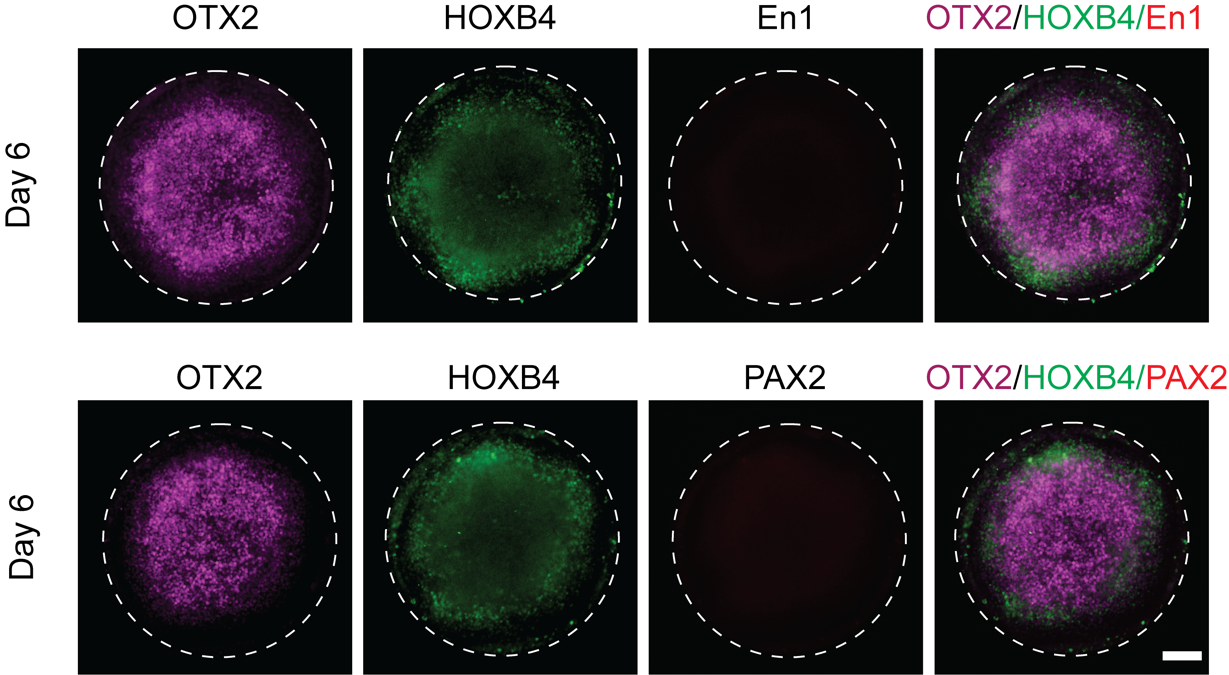

**Figure S6. Characterization of cell fates in self-organized anteroposterior patterned human neural tissue**. Representative immunostaining fluorescence images showing the spatial expression of neural markers: OTX2 (forebrain and midbrain marker), HOXB4 (hindbrain marker), EN1 (midbrain and hindbrain marker), and PAX2 (midbrain and hindbrain marker). In this experiment, H9 human ESCs were differentiated with the induction protocol and immunostained on day 6. Scale bar, 100 μm.

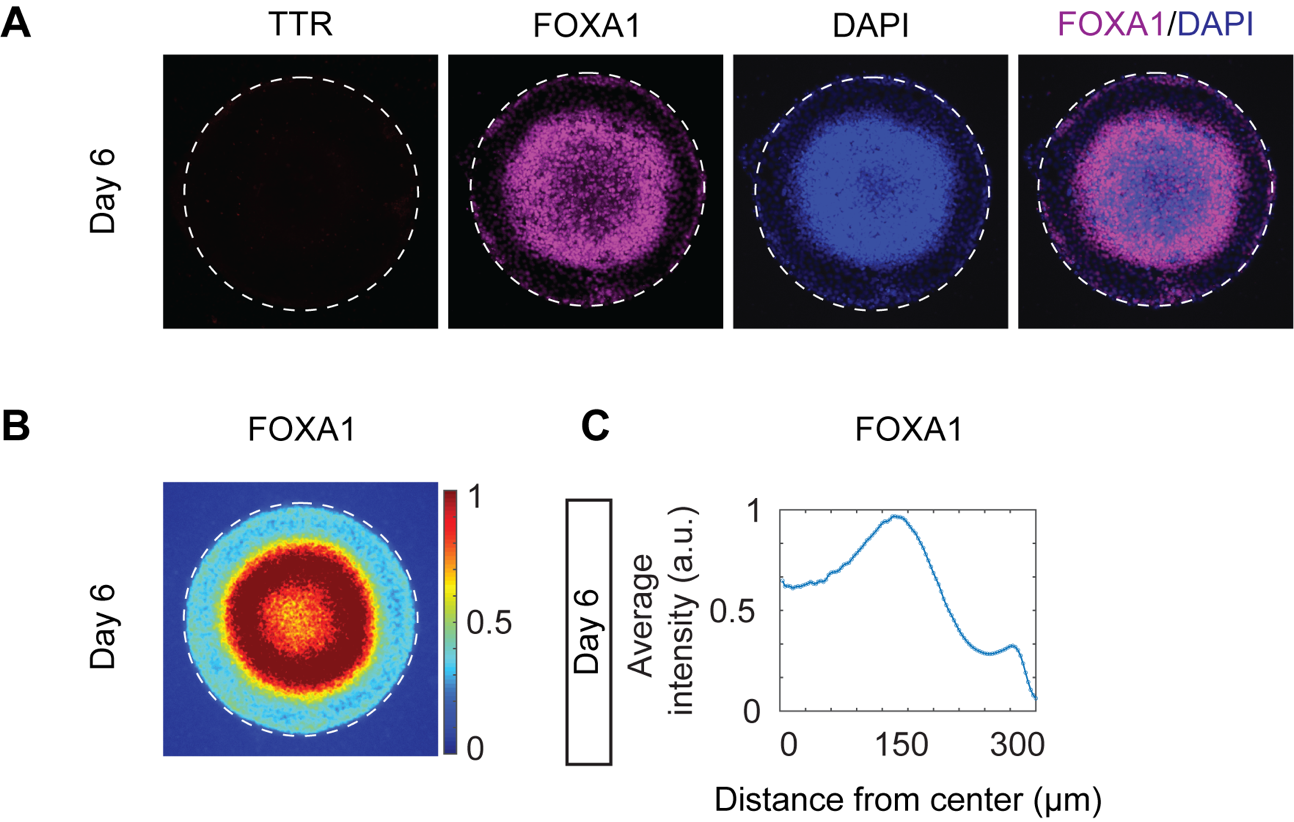

**Figure S7. Further characterization of cell fates in the micropattern with immunofluorescence staining**. (A). Representative immunostaining fluorescence images showing the spatial expression of FOXA1 (a marker for ventral midbrain dopaminergic progenitors) and TTR (a choroid plexus marker) on day 6. Cell nuclei were counterstained with DAPI. (B). Colorimetric maps showing the average fluorescent intensity of FOXA1 staining. The intensity of each pixel of the images was normalized to the maximum value for each image. n > 15. (C). Plots showing the quantitative average intensity of FOXA1 in relation to the distance to the center of the pattern. In this experiment, H9 human ESCs were differentiated with the induction protocol and immunostained on day 6. Scale bar, 100 μm.

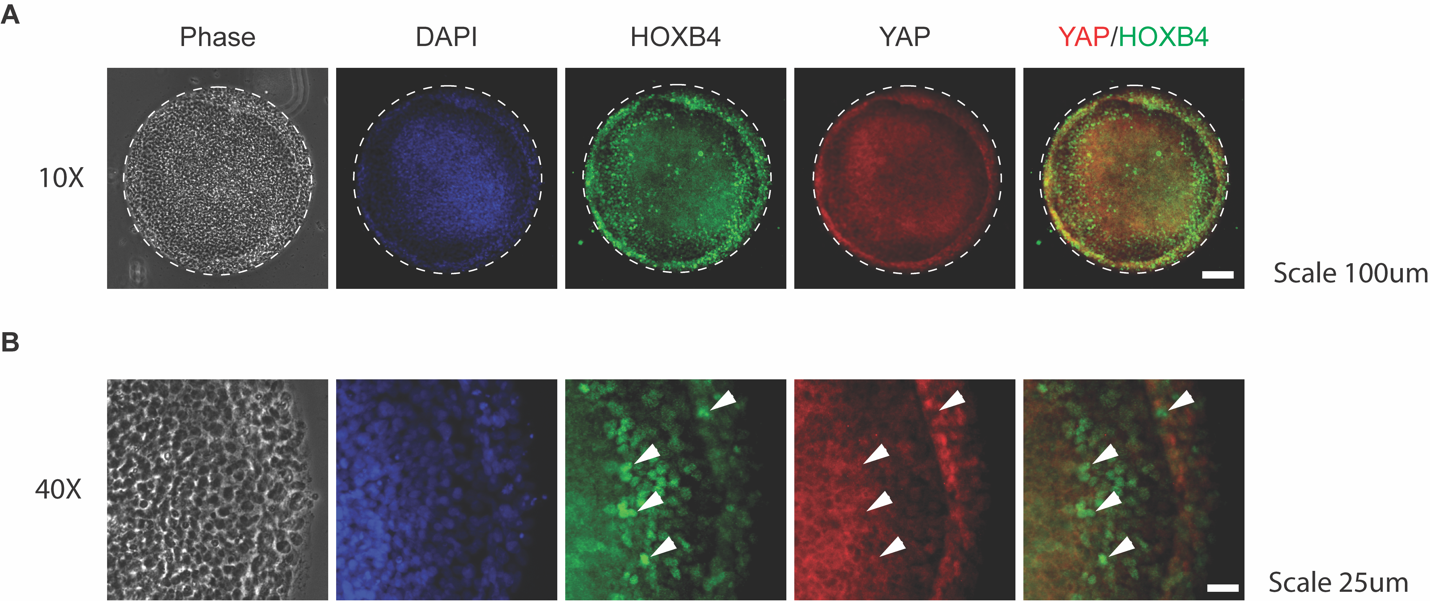

**Figure S8. The co-localization of HOXB4 and nuclei YAP was minimal.** (A). Representative phase-contrast and immunostaining fluorescence images showing the expression of hindbrain markers HOXB4 and the localization of YAP on day 6. 10× objective was used. The YAP and HOXB4 merged image was shown. Cell nuclei were counterstained with DAPI. Scale bar, 100 μm. (B). Representative phase-contrast and immunostaining fluorescence images showing the expression of hindbrain markers HOXB4 and the localization of YAP on day 6. 40× objective was used. The YAP and HOXB4 merged image was shown. Some representative HOXB4+ cells with cytoplasmic YAP were marked with arrowheads. Cell nuclei were counterstained with DAPI. Scale bar, 25 μm.

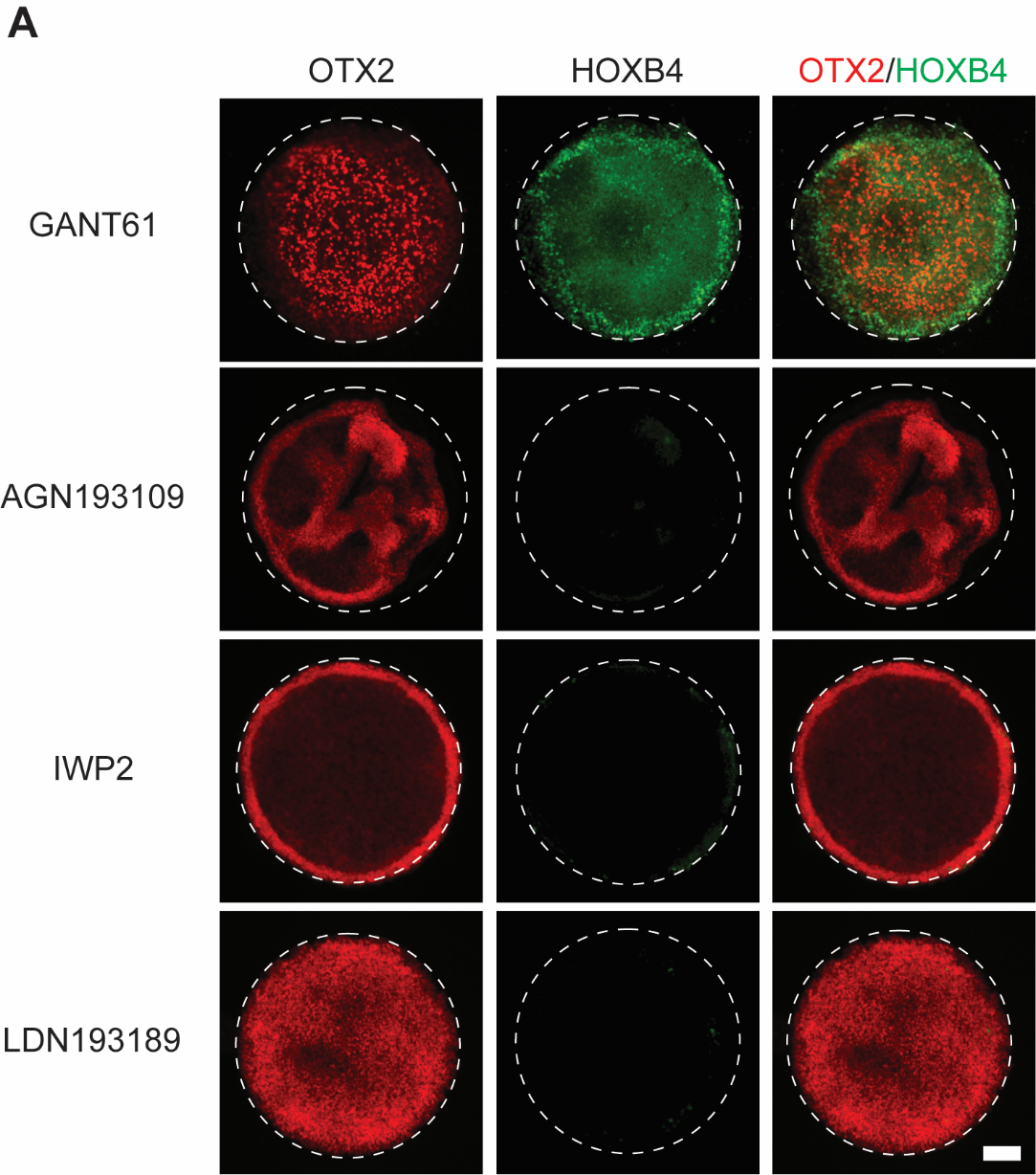

**Figure S9. The effects of SHH, RA, WNT, or BMP signals inhibitors treatment on midbrain/hindbrain patterning.** Representative immunostaining fluorescence images showing the spatial expression of OTX2 and HOXB4 of human ESCs treated with GANT61 (a sonic hedgehog signaling inhibitor, 10 µM), AGN193109 (a retinoic acid signaling inhibitor, 100 nM), IWP2 (a WNT signaling inhibitor, 3 µM), and LDN193189 (a BMP signaling inhibitor, 500 nM) on day 6. Scale bar, 100 μm.

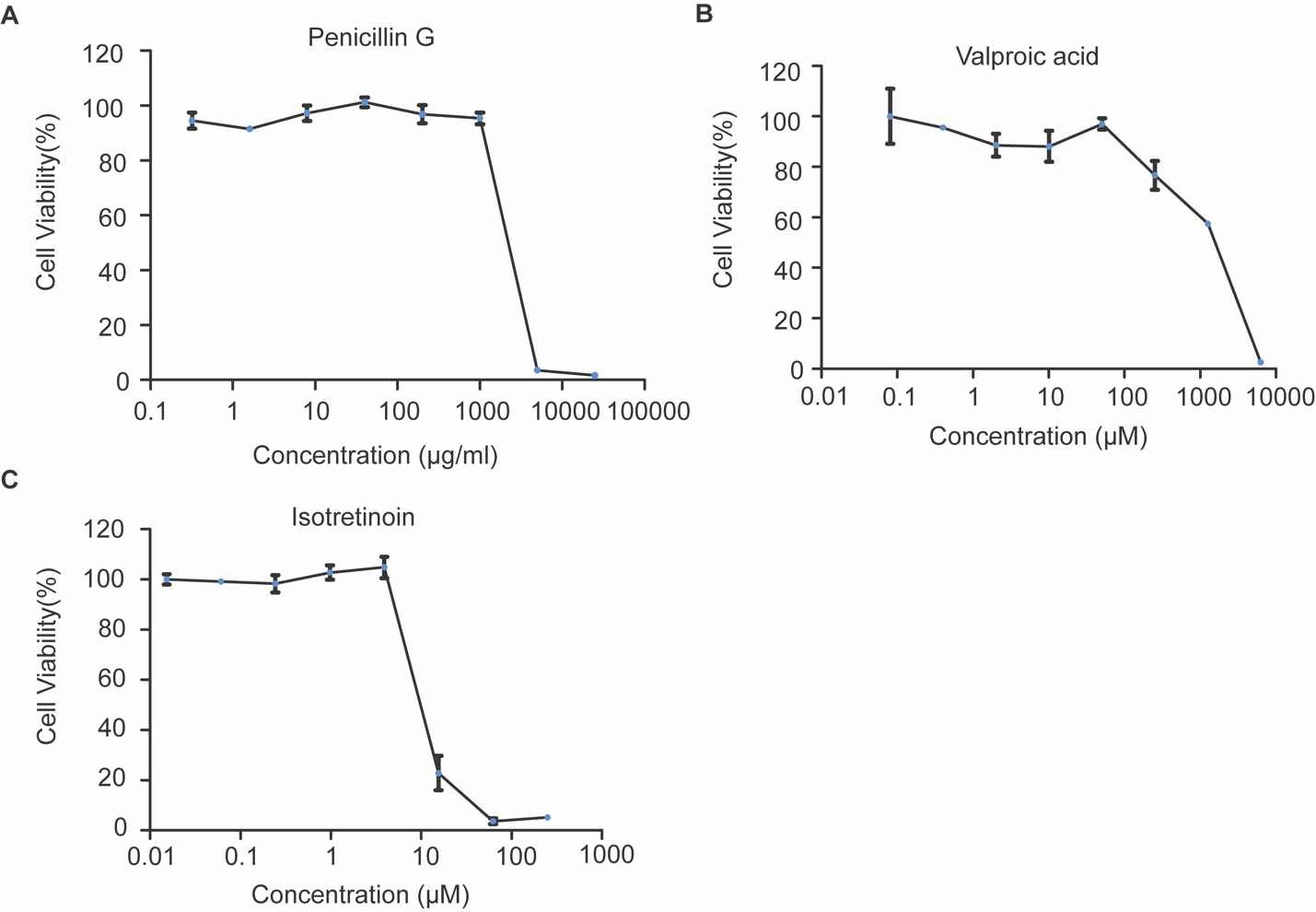

**Figure S10. Cell viability curves for penicillin G, VPA, and isotretinoin treatment in H9 human embryonic stem cells.** (A). H9 cells were treated with 8 different concentrations of a non-teratogenic drug, penicillin G (n = 2) for two days. (B). H9 cells were treated with 8 different concentrations of valproic acid (VPA, n = 2) for two days. (C). H9 cells were treated with 8 different concentrations of isotretinoin (n = 3) for two days.

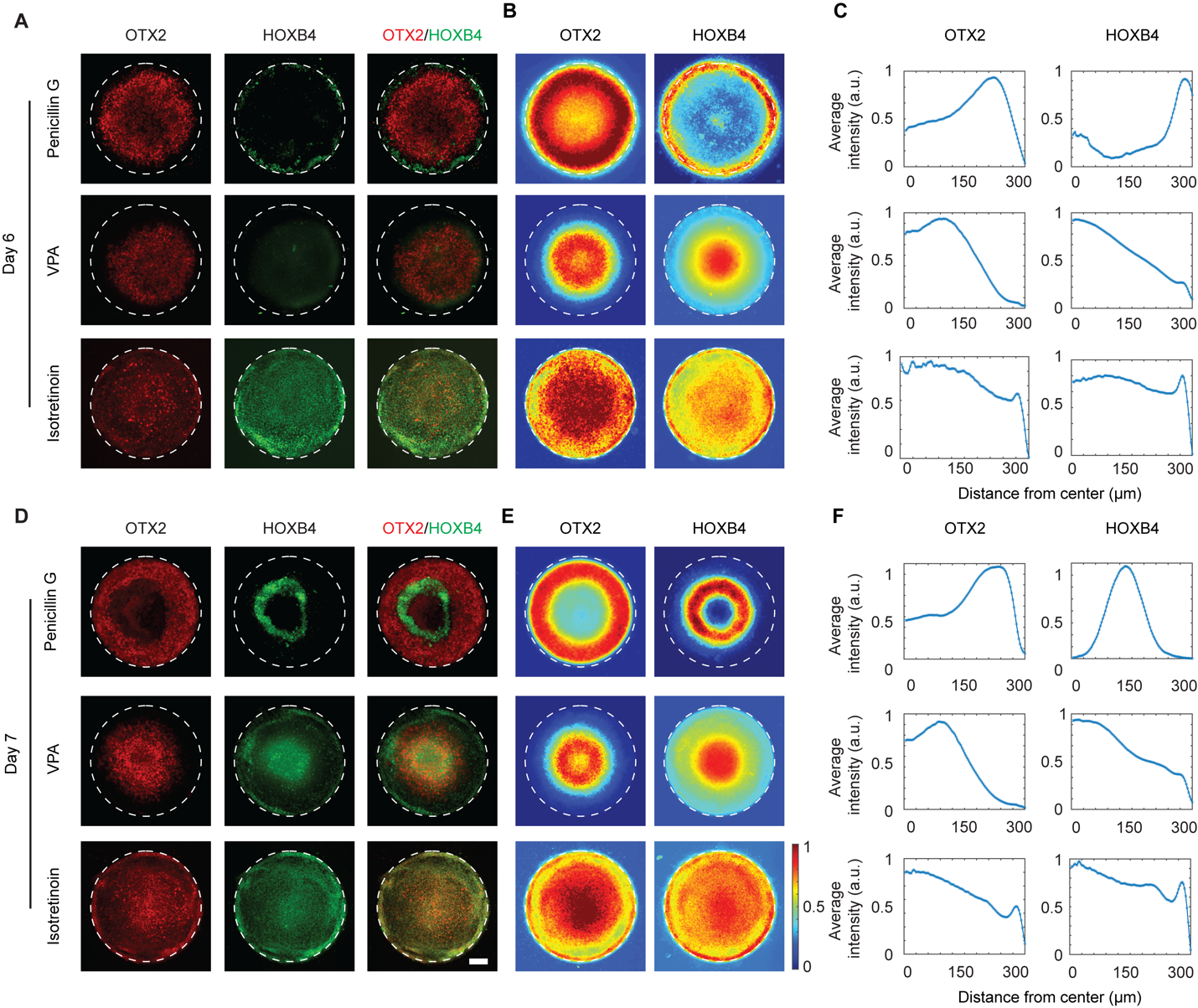

**Figure S11. The effects of low dosages of penicillin G, valproic acid, and isotretinoin on anteroposterior patterning of OTX2 and HOXB4.** (A). Representative immunostaining fluorescence images showing the spatial expression of OTX2 and HOXB4 in induced human ESCs treated with low dosages of penicillin G, valproic acid, and isotretinoin on day 6. (B). Colorimetric maps showing the average fluorescent intensity of OTX2 and HOXB4 staining. The intensity of each pixel of the images was normalized to the maximum value for each image. n > 15. (C). Plots showing the quantitative average intensity in relation to the distance to the center of the pattern. (D). Representative immunostaining fluorescence images showing the spatial expression of OTX2 and HOXB4 in induced human ESCs treated with low dosages of penicillin G, valproic acid, and isotretinoin on day 7. (E). Colorimetric maps showing the average fluorescent intensity of OTX2 and HOXB4 staining. The intensity of each pixel of the images was normalized to the maximum value for each image. n > 15. (F). Plots showing the quantitative average intensity in relation to the distance to the center of the pattern. (G). The percentage of area covered by OTX2+ midbrain in DMSO control, penicillin G, VPA, and isotretinoin treated groups. Scale bar, 100 μm. Data are represented as mean ± s.e.m. ***, P < 0.001.

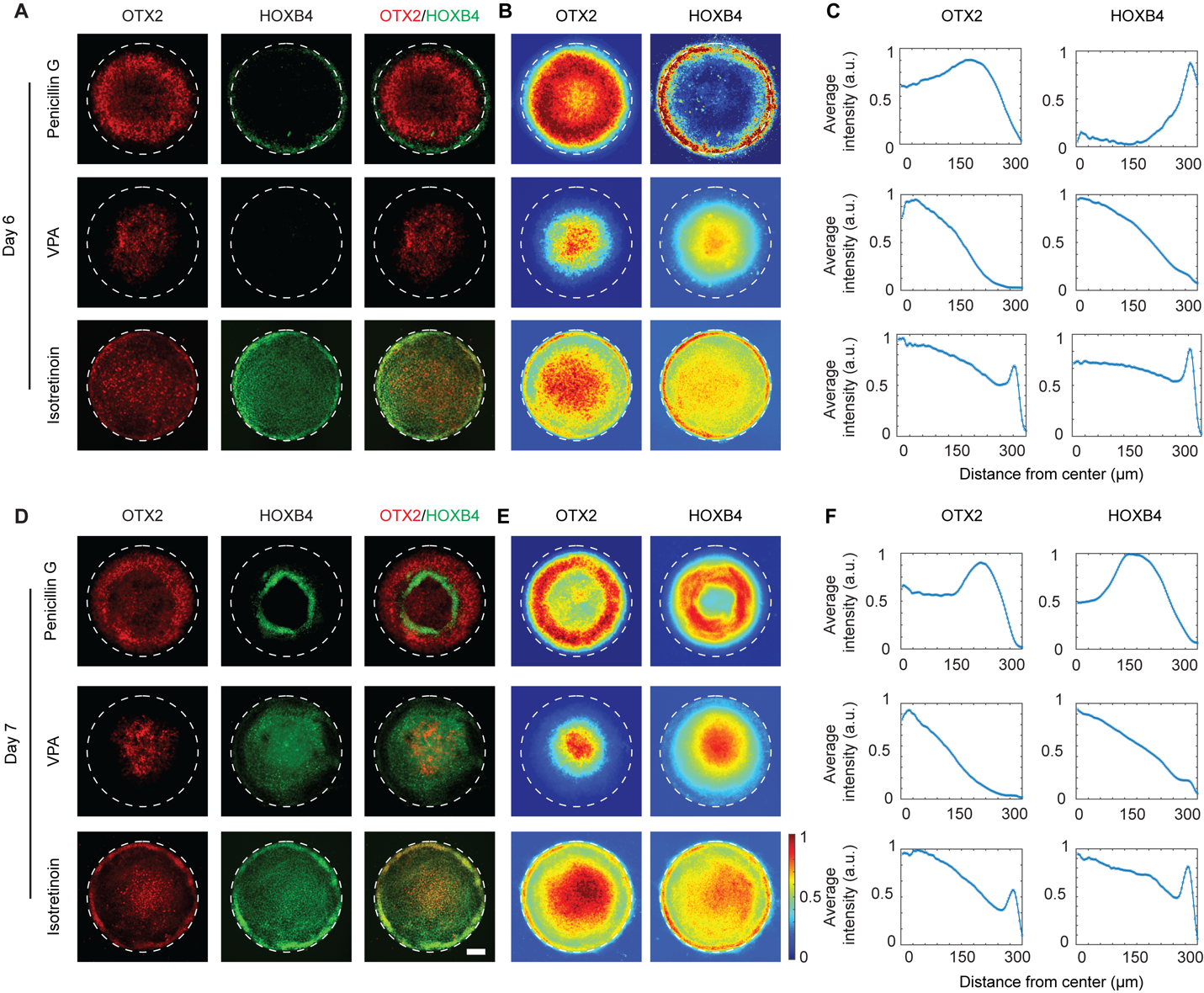

**Figure S12. The effects of high dosages of penicillin G, valproic acid, and isotretinoin on anteroposterior patterning of OTX2 and HOXB4.** (A). Representative immunostaining fluorescence images showing the spatial expression of OTX2 and HOXB4 in induced human ESCs treated with high dosages of penicillin G, valproic acid, and isotretinoin on day 6. (B). Colorimetric maps showing the average fluorescent intensity of OTX2 and HOXB4 staining. The intensity of each pixel of the images was normalized to the maximum value for each image. n > 15. (C). Plots showing the quantitative average intensity in relation to the distance to the center of the pattern. (D). Representative immunostaining fluorescence images showing the spatial expression of OTX2 and HOXB4 in induced human ESCs treated with high dosages of penicillin G, valproic acid, and isotretinoin on day 7. (E). Colorimetric maps showing the average fluorescent intensity of OTX2 and HOXB4 staining. The intensity of each pixel of the images was normalized to the maximum value for each image. n > 15. (F). Plots showing the quantitative average intensity in relation to the distance to the center of the pattern. (G). The percentage of area covered by OTX2+ midbrain in DMSO control, penicillin G, VPA, and isotretinoin treated groups. Scale bar, 100 μm. Data are represented as mean ± s.e.m. ***, P < 0.001.

Table S1. List of antibodies used in immunofluorescence

| **Protein** | **Vendor** | **Catalog number** | **RRID** | **Dilution** |
| --- | --- | --- | --- | --- |
| PAX6 | Abcam | Ab78545 | n/a | 1:100 |
| N-cadherin | Proteintech | 66219-1-Ig | AB_2881610 | 1:100 |
| HoxB4 | DSHB | I12 | AB_2119288 | 1:20 |
| OTX2 | R&D Systems | AF1979 | n/a | 1:100 |
| EN1 | DSHB | 4G11 | AB_2314371 | 1:20 |
| PAX2 | DSHB | 1A7 | AB_2722284 | 1:10 |
| FOXG1 | Clontech | M227 | n/a | 1:200 |
| GBX2 | Proteintech | 21639-1-AP | n/a | 1:200 |
| YAP | Santa Cruz | Sc-101199 | n/a | 1:100 |
| FOXA1 | Santa Cruz | Sc-514695 | n/a | 1:100 |
| TTR | Bio-Rad | AHP1837 | n/a | 1:200 |

Table S2. List of compounds for drug screening, their IC_25_, and their peak plasma concentrations (C_max_) *in vivo*.

| **Compounds** | **IC_25_** | **C_max_** | **Reference for C_max_** |
| --- | --- | --- | --- |
| Penicillin G | 1946 µg/ml | 400 µg/ml | Plaut et al. 1969 (*69*) |
| VPA | 0.74 mM | 0.574 mM | Reed et al. 2006 (70) |
| Isotretinoin | 3.39 µM | 2.87 µM | Roche. Accutan® (isotretinoin) [package insert]. 2010 (71) |

Table S3. List of compounds for drug screening and the selected high, medium, and low concentrations for testing.

| **Compounds** | **High** | **Medium** | **Low** |
| --- | --- | --- | --- |
| Penicillin G | 1000 µg/ml | 200 µg/ml | 40 µg/ml |
| VPA | 0.25 mM | 0.12 mM | 0.06 mM |
| Isotretinoin | 3.25 µM | 1.63 µM | 1.08 µM |

**Movie S1. The morphologic change of micropatterned cells from day 6 to day 7.** Micropatterned human embryonic cells were differentiated into midbrain and hindbrain cell progenitors. The cells were folded inwardly and then formed a 3D multilayer structure in 3D from day 6 to day 7. Images were taken every one hour. The play speed of the video is two frames per second. Scale bar, 100 µm.
